# POT1 recruits and regulates CST–Polα/Primase at human telomeres

**DOI:** 10.1101/2023.05.08.539880

**Authors:** Sarah W. Cai, Hiroyuki Takai, Thomas Walz, Titia de Lange

**Affiliations:** Laboratory of Cell Biology and Genetics, The Rockefeller University; New York, NY, USA; Laboratory of Molecular Electron Microscopy, The Rockefeller University; New York, NY, USA

**Keywords:** telomere, DNA replication, CST, Polymerase α/Primase, shelterin, POT1, cryo-EM, DNA-protein complex, phospho-regulation

## Abstract

Telomere maintenance requires extension of the G-rich telomeric repeat strand by telomerase and fill-in synthesis of the C-rich strand by Polα/Primase. Telomeric Polα/Primase is bound to Ctc1-Stn1-Ten1 (CST), a single-stranded DNA-binding complex. Like mutations in telomerase, mutations affecting CST–Polα/Primase result in pathological telomere shortening and cause a telomere biology disorder, Coats plus (CP). We determined cryogenic electron microscopy structures of human CST bound to the shelterin heterodimer POT1/TPP1 that reveal how CST is recruited to telomeres by POT1. Phosphorylation of POT1 is required for CST recruitment, and the complex is formed through conserved interactions involving several residues mutated in CP. Our structural and biochemical data suggest that phosphorylated POT1 holds CST–Polα/Primase in an inactive auto-inhibited state until telomerase has extended the telomere ends. We propose that dephosphorylation of POT1 releases CST–Polα/Primase into an active state that completes telomere replication through fill-in synthesis.

## Introduction

Telomeres prevent the inappropriate activation of the DNA damage response at the ends of linear chromosomes. Human telomeric DNA is comprised of double-stranded (ds) 5’-TTAGGG-3’ repeats that terminate in a single-stranded (ss) 3’ overhang. This DNA protects chromosome ends through its interaction with the six-subunit shelterin complex (reviewed in ^1^). With each cell cycle, incomplete DNA synthesis and nucleolytic resection at telomere ends results in attrition of the telomeric DNA that can lead to excessive telomere shortening if unchecked. Telomere maintenance is critical for the long-term proliferation of human cells and involves two distinct DNA synthesizing enzymes: telomerase, which elongates the G-rich strand, and CST–Polα/Primase, which maintains the C-rich repeats through fill-in synthesis (reviewed in ^2^ and (Cai and de Lange, in revision)).

Telomere extension by telomerase takes place after DNA replication and requires exonucleolytic processing of the leading-strand DNA synthesis product to create a 3’ overhang that serves as the telomerase primer (reviewed in ^3^). Telomerase is recruited to telomeres by the TPP1 subunit of shelterin, whose TEL patch directly interacts with the reverse transcriptase hTERT ^4–7^. The extension of telomeres by telomerase is regulated by several shelterin components, which appear to maintain telomere length homeostasis through a cis-acting negative feedback loop ^8–10^. This telomere length homeostasis pathway was recently shown to be critical for tumor suppression through telomere shortening, which prevents a multitude of cancer types in the human species ^11,12^. Whereas much is known about how telomerase acts at telomere ends ^13^, the mechanism of CST–Polα/Primase recruitment, regulation, and action is not well understood.

CST is a heterotrimeric ssDNA-binding protein that shares structural homology with the major eukaryotic ssDNA-binding protein, Replication Protein A (RPA) ^14–16^. However, it is likely that CST evolved independently in an archaeal ancestor and became specialized for telomere maintenance in eukaryotes (^17^; reviewed in (Cai and de Lange, in revision)). In metazoans, the large Ctc1 subunit of CST contains two sets of tandem oligosaccharide/oligonucleotide-binding (OB) folds ^16^ that are connected by a three-helix bundle, termed the Acidic Rpa1 OB-binding Domain-like three-helix bundle (Ctc1^ARODL^) based on similarity to its archaeal counterpart ^17^. Ctc1^OB-A/B/C/D^ and the Ctc1^ARODL^ comprise the metazoan-specific N-terminal domain of the protein, whereas Ctc1^OB-E/F/G^ are ancestral and mediate ssDNA binding and trimerization (reviewed in (Cai and de Lange, in revision)). Stn1, comprising an N-terminal OB fold (Stn1^N^) and C-terminal tandem winged helix-turn-helix (wHTH) domains (Stn1^C^), bridges Ten1’s single OB fold to Ctc1 ^17^.

Recent cryo-EM structures of CST–Polα/Primase revealed how CST interacts with Polα/Primase in two distinct conformations. Polα/Primase can adopt a compact, auto-inhibited state ^18^ or an extended active state ^19^. CST binds the auto-inhibited Polα/Primase using the Ctc1 N-terminal domain, and this interaction is conserved in metazoans ^18^. This inactive complex was proposed to represent a Recruitment Complex (RC) that describes the conformation of CST–Polα/Primase as it is brought to the telomere prior to fill-in. Human CST–Polα/Primase was also captured in an active Pre-Initiation Complex (PIC), in which CST uses its ancestral regions (Ctc1^OB-E/F/G^, Stn1, and Ten1) to stabilize and stimulate the extended Polα/Primase ^19^. A significant conformational change is required to transition from the RC to PIC, but how this change is regulated is unclear.

The importance of CST–Polα/Primase in the maintenance of telomeres is illustrated by CP, a rare pediatric condition characterized by the ophthalmological disorder Coats disease *plus* additional associated systemic disorders primarily affecting the brain, bone, and gastrointestinal tract ^20–29^. CP is caused by hypomorphic mutations in Ctc1, Stn1, or POT1 that reveal their phenotype only when the second copy of the gene is non-functional. Complete loss of CST leads to shortening of the C-rich strand due to incomplete lagging-strand DNA synthesis at the telomere ends (Takai, Aria, Yeeles, and de Lange, in preparation) and nucleolytic resection of the C-strand at leading-end telomeres ^30–33^.

It has remained unclear how CST–Polα/Primase is recruited to telomeres. Initial experiments in the mouse showed that CST interacts with one of the two mouse POT1 proteins, mPOT1b ^30^. Mutations in POT1b diminish CST recruitment and lead to lack of C-strand fill-in at telomeres. However, human POT1, which combines the functions of mouse mPOT1a and mPOT1b in a single protein ^34^, does not form a complex with CST detectable in co-IP experiments (Fig. 1A). In contrast, human TPP1 readily co-IPs with CST and was therefore thought to play a primary role in recruitment of human CST ^35^. Yet, both POT1 and TPP1 were reported to interact with Ctc1 in one yeast two-hybrid assay ^36^ but not in another ^15^. Another puzzling observation is presented by the CP POT1 mutation (S322L), which recapitulates the phenotypes of diminished CST function ^27^.

**Fig. 1.**
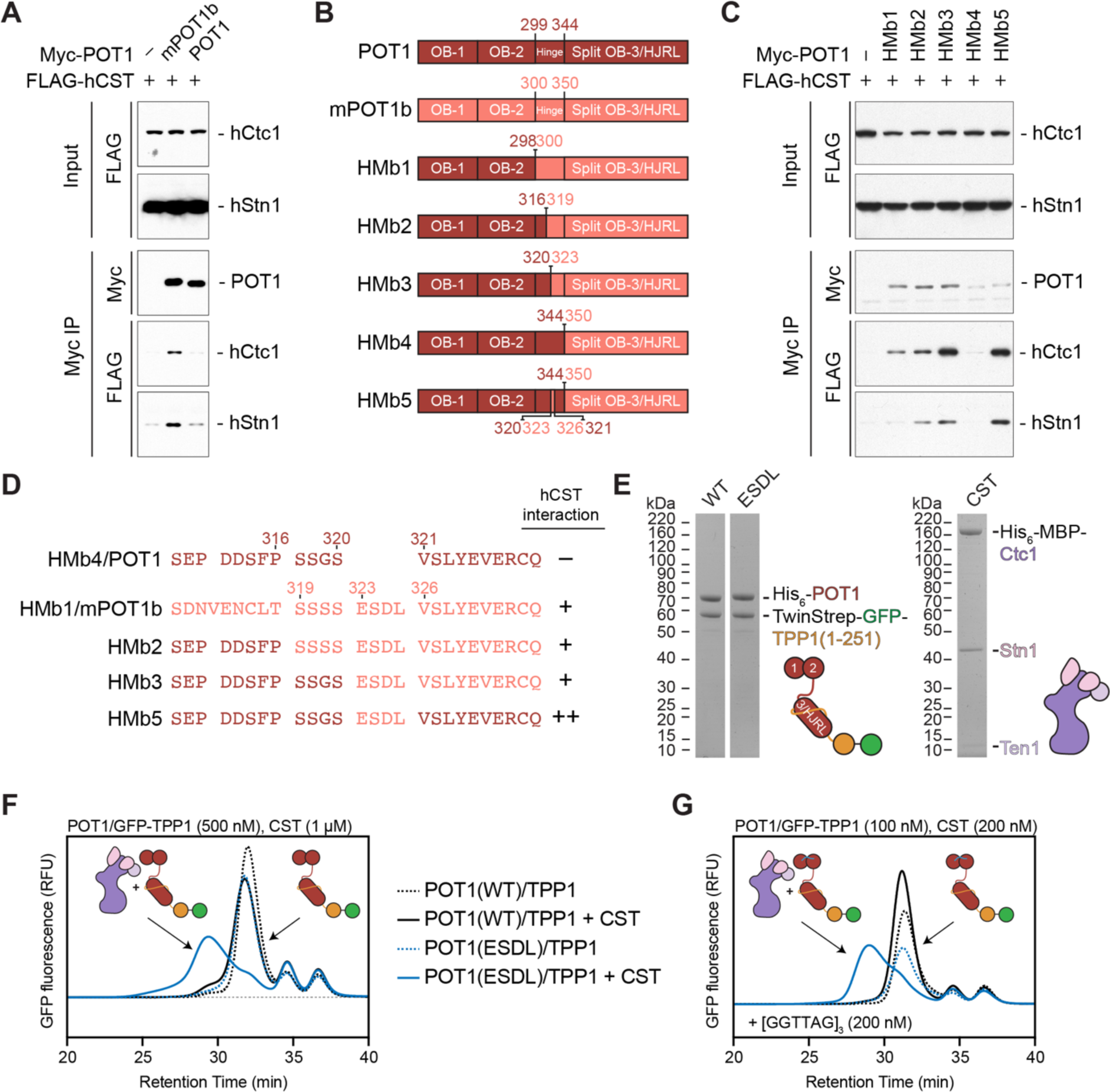
POT1-CST interaction. (**A**) Immunoblot of anti-Myc co-IPs of Myc-tagged POT1 proteins and FLAG-tagged CST from co-transfected 293T cells showing that mPOT1b interacts with human CST better than human POT1. Immunoblots were probed with anti-Myc and anti-FLAG antibodies. (**B**) Design of constructs encoding chimeric swaps between mPOT1b (red) and human POT1 (salmon). HMb: **H**uman/**M**ouse POT1**b** swap; OB: oligosaccharide/oligonucleotide-binding domain; HJRL: Holliday junction resolvase-like domain. (**C**) Immunoblots of CST binding by chimeric human-mouse POT1 proteins. Anti-Myc co-IPs of Myc-tagged POT1 constructs, and FLAG-tagged CST from co-transfected 293T cells showing that aa 323-326 of mPOT1b are important for the CST interaction (comparison of HMb4 and HMb5). Immunoblots were probed with anti-Myc and anti-FLAG antibodies. (**D**) Sequence alignment of a region in the POT1 hinge (aa 309-330) (left) and the effect of the sequence on the interaction with CST (right). HMb4 contains the POT1 hinge and HMb1 contains the mPOT1b hinge. HMb3 and HMb5 contain the same hinge sequence shown, but they represent different constructs as shown in (**B**). (**E**) Cartoon schematics and gels of purified POT1/TPP1 and CST proteins used for fluorescent size-exclusion chromatography (FSEC) analysis (**F-G**) FSEC analysis of the interaction between CST and POT1(WT)/GFP-TPP1 (blue) or POT1(ESDL)/GFP-TPP1 (red) in the absence (**F**) or presence (**G**) of telomeric ssDNA. Traces without CST are shown as dashed lines and color key is the same for both panels. RFU: relative fluorescence units.

Here, we present structural and biochemical data revealing that human CST primarily interacts with POT1. This interaction is dependent on POT1 phosphorylation, reconciling the negative co-IP data. We identified four amino acids in mPOT1b that, when inserted into the human protein, result in POT1 phosphorylation in both human cells and insect cells. With the phosphorylated form of POT1, we reconstituted CST–POT1/TPP1 complexes in the presence and absence of telomeric ssDNA and determined their structures using cryo-EM. POT1/TPP1 has highly flexible regions that previously prevented high-resolution structure determination ^37,38^, but in our structures, POT1 is held in a single conformation and interacts with Ctc1 at three distinct sites. The structure indicates that phosphorylation drives charge complementarity between the two proteins. Two other interfaces are independent of the mPOT1b insertion and confirm that human POT1 is the primary recruiter of CST. Several CP mutations map to or near the primary interface between Ctc1^OB-D^, Ctc1^ARODL^, and the POT1 C-terminal half. On the other end, the POT1 N-terminal OB folds interact with Ctc1’s ssDNA-binding site. This interaction directly competes with CST ssDNA binding and Polα/Primase binding in the active PIC conformation, whereas the major interface on Ctc1 for Polα/Primase binding in the auto-inhibited RC conformation is unobstructed. Together, our data identify a phosphorylation-dependent switch in POT1 that controls its interaction with CST and can regulate the activity of CST–Polα/Primase at the telomere. This provides a molecular basis for how shelterin recruits and regulates CST–Polα/Primase and reveals the mechanism of mutations in CST and POT1 that cause telomere dysfunction in CP.

## Results

### Reconstitution of a stable CST–POT1/TPP1 complex

To understand how POT1/TPP1 recruits CST, we first examined how TPP1, the established CST interactor, binds. Co-IP data indicated that a region within the C-terminal 20 residues of TPP1 is necessary for its interaction with CST (Fig. S1A). This region of TPP1 constitutes its TIN2-interaction domain (TPP1^TID^) and is required to anchor the POT1/TPP1 heterodimer to the duplex telomeric DNA-binding proteins in shelterin ^39^. Indeed, co-IP experiments showed that TIN2 competes with CST for TPP1 binding (Fig. S1B). AlphaFold-Multimer ^40^ models of TPP1^TID^ bound to Ctc1 predicted that TPP1^TID^ binds to Ctc1^OB-B^ and uses the same amino acids used for binding to TIN2, suggesting that TPP1 binds either TIN2 or Ctc1 but not both at the same time (Fig. S1C-D). Since its association with TIN2 is required for TPP1/POT1 to function at telomeres ^41,42^, we consider it unlikely that the TPP1^TID^ is involved in the recruitment of CST. As predicted, a recent report confirmed that loss of the interaction between TPP1^TID^ and Ctc1^OB-B^ did not abolish fill-in by CST–Polα/Primase ^43^. We therefore omitted the TPP1^TID^ from our analysis to limit heterogeneity from the disordered TPP1 C-terminus (aa 252-458).

Interestingly, although human POT1 does not form a stable complex with CST in co-IP experiments, mPOT1b readily bound to human CST (Fig. 1A). This finding suggested to us that mPOT1b can constitutively bind CST whereas the binding of human POT1 to CST may be regulated, perhaps in a cell cycle-dependent manner. To determine which part of mPOT1b mediates the CST interaction, we generated chimeric proteins encoding swaps between mPOT1b and human POT1 (Fig. 1B). When swapped with the human POT1 C-terminus (aa 320-634), the mPOT1b C-terminus (aa 323-640) was sufficient to mediate an interaction with CST (Fig. 1B-C). This interaction was further narrowed down to the C-terminal half of the mPOT1b hinge (aa 323-350) (Fig. 1B-C). mPOT1b contains four additional residues in this region that were a key determinant of the interaction (ESDL, aa 323-326, Fig. 1B-D). When inserted into the human POT1 hinge (between aa 320-321, Fig. 1D), there was a robust interaction between CST and POT1(ESDL)/TPP1 compared to wild-type POT1/TPP1 *in vitro* (Fig. 1E-G). POT1(ESDL)/TPP1 formed a complex with CST in the presence or absence of ssDNA, but addition of a telomeric [GGTTAG]_3_ ssDNA allowed complex formation at lower protein concentrations (Fig. 1F-G; Fig. S2A).

We reconstituted two complexes of CST–POT1(ESDL)/TPP1. With the addition of the 18-nt [GGTTAG]_3_ ssDNA, we could reconstitute a stable complex containing full-length CST, full-length POT1(ESDL), and TPP1^OB+RD^ (aa 1-251; Fig. 2A; Fig. S2B). Without ssDNA, the complex dissociated at the low concentrations (< 200 μM) used for negative-stain and cryo-EM (Fig. S2A). To increase the local concentration of CST relative to POT1(ESDL)/TPP1 for structural studies, we fused aa 1-403 of TPP1 to the N-terminus of Stn1, retaining 162 aa of TPP1’s disordered serine-rich linker to allow for flexibility between the two polypeptides (Fig. 2A; Fig. S2C). This fusion allowed for purification of an apo CST–POT1(ESDL)/TPP1 complex in the absence of ssDNA (Fig. S2D).

**Fig. 2.**
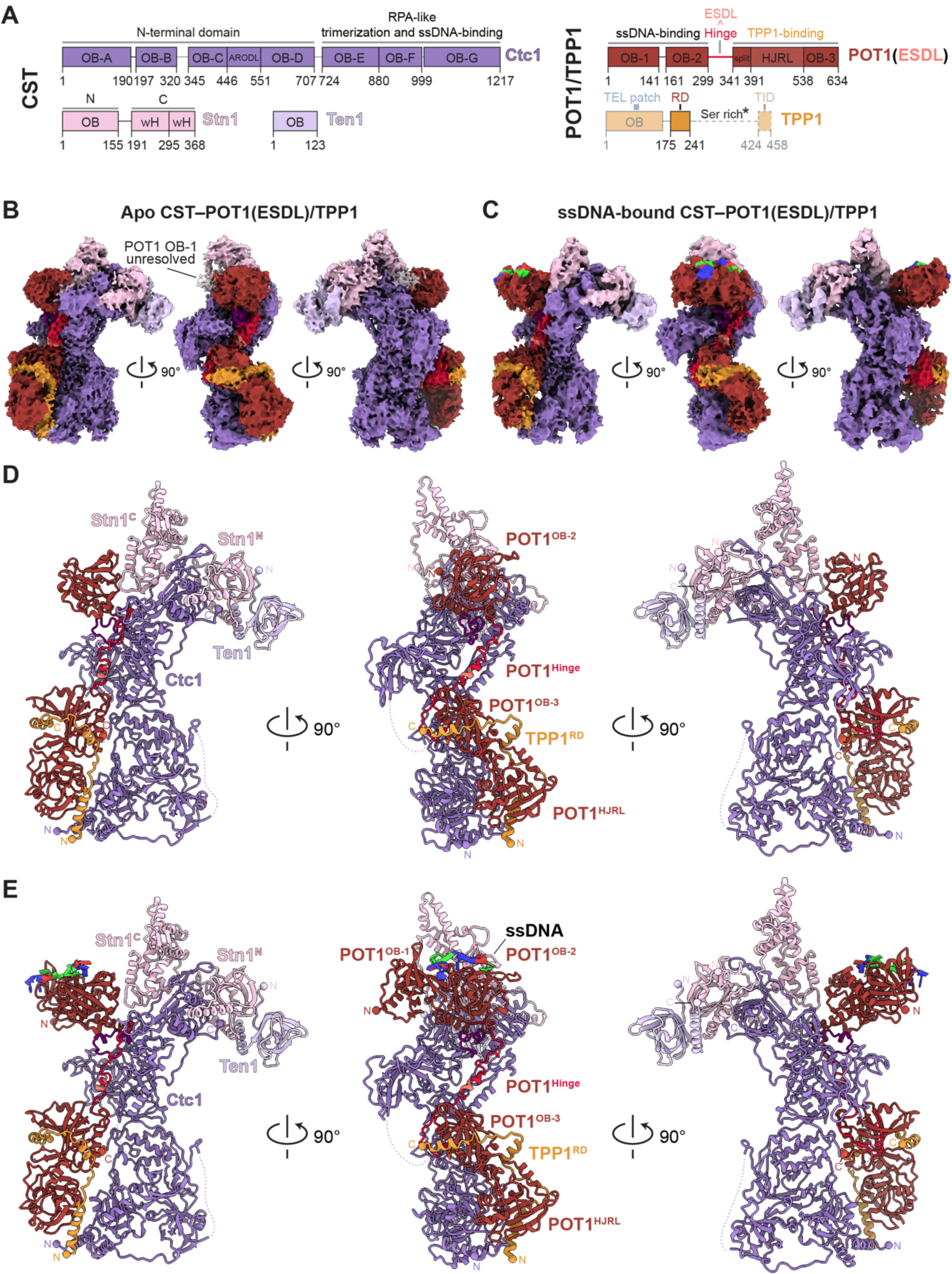
Cryo-EM structures of CST–POT1(ESDL)/TPP1 complexes. (**A**) Domain organization of CST and the shelterin subunits POT1(ESDL) and TPP1. Regions not observed in the cryo-EM maps are shown at 50% opacity. Dashed lines indicate regions not included in the expression construct. *The TPP1 serine-rich linker was included in the apo complex but not the ssDNA-bound complex. ARODL: Acidic Rpa1 OB-binding Domain-like 3-helix bundle; wH: winged helix-turn-helix domain; RD: recruitment domain, TID: TIN2-interacting domain. For other abbreviations see Fig. 1. (**B**) Cryo-EM reconstruction of apo CST–POT1(ESDL)/TPP1 at 3.9-Å resolution (see also Fig. S3). (**C**) Cryo-EM reconstruction of ssDNA-bound CST–POT1(ESDL)/TPP1 at 4.3-Å resolution (see also Fig. S4) (**D**-**E**) Atomic models for the apo and ssDNA-bound CST–POT1(ESDL)/TPP1 complexes, respectively. See also Video S1.

### Cryo-EM structures of CST–POT1/TPP1

Negative-stain EM analysis showed additional density attributable to the addition of POT1(ESDL)/TPP1 in the two complexes (Fig. S2E). We determined cryo-EM structures of apo and ssDNA-bound CST–POT1(ESDL)/TPP1 at overall resolutions of 3.9- and 4.3-Å, respectively (Fig. 2; Fig. S3-4; Video S1; Table 1). Into these density maps, we could unambiguously place models of the individual subunits predicted by AlphaFold2 ^44,45^ or previously determined by X-ray crystallography or cryo-EM as a starting point to build the models (Table 1). The map quality of the apo CST–POT1(ESDL)/TPP1 reconstruction was better (Fig. S3H), so we primarily used that map for model building. The resulting model from the apo complex was then used as a starting model to build the ssDNA-bound complex. We observed continuous density for the POT1 flexible hinge and were able to build its Cα backbone *de novo*, allowing us to generate an experimental structure of full-length POT1(ESDL) (Fig. 2D-E, Fig. S3-4; Video S1). The POT1 C-terminus, consisting of POT1^OB-3^ and POT1^HJRL^ (Fig. 2A), is bound by TPP1’s recruitment domain (TPP1^RD^), as observed in previous crystal structures (PDB 5H65 ^46^ and PDB 5UN7 ^47^) (Fig. 2D-E). TPP1^RD^ does not appear to interact directly with CST in either structure. TPP1’s N-terminal OB-fold (TPP1^OB^) was not resolved, presumably due to flexible attachment. At low contouring thresholds, a cylindrical density was observed that likely represents TPP1^OB^ (Fig. S5B). This putative position of TPP1^OB^ suggested that it is not bound to CST, and TPP1^OB^ was indeed dispensable in the interaction of POT1(ESDL)/TPP1 with CST (Fig. S5C).

**Table 1.** Cryo-EM data collection, refinement, and validation statistics.

|  |  |  |
| --- | --- | --- |
| Complex | apo CST–POT1/TPP1 | ssDNA–CST–POT1/TPP1 |
| EMDB ID | 40659 | 40660 |
| PDB ID | 8SOJ | 8SOK |
| **Data collection** |  |  |
| Microscope | Titan Krios | Titan Krios |
| Detector | K3 | K3 |
| Voltage (kV) | 300 | 300 |
| Electron exposure (e<sup>-</sup>/Å<sup>2</sup>) | 51 | 51 |
| Magnification (kx) | 105 | 105 |
| Pixel size (Å) | 0.86 | 0.86 |
| Defocus range (μm) | -1 to -2.5 | -1 to -2.5 |
| Micrographs collected | 15,201 | 31,914 |
| **Reconstruction** |  |  |
| Symmetry imposed | C1 | C1 |
| Initial particle images (No.) | 4,835,578 | 2,213,253 |
| Final particle images (No.) | 132,356 | 76,359 |
| Pixel size (Å) | 0.86 | 0.86 |
| Box size (px) | 352 | 352 |
| Map resolution (Å) | 3.9 | 4.3 |
| FSC threshold | 0.143 | 0.143 |
| Map sharpening B factor (Å<sup>2</sup>) | -130 | -150 |
| **Model composition** |  |  |
| Non-hydrogen atoms | 17,227 | 18,574 |
| Protein residues | 2195 | 2335 |
| Nucleotides | 0 | 10 |
| Ligands | 0 | 0 |
| Metals | 2 (Zn<sup>2+</sup>) | 2 (Zn<sup>2+</sup>) |
| **Refinement** |  |  |
| Initial models used (PDB code) | 6W6W, 5H65, 1XJV, AF-Q2NKJ3 (AlphaFold database) |  |
| Model-to-map CC (mask) | 0.77 | 0.67 |
| Model-to-map CC (volume) | 0.74 | 0.64 |
| B factors (Å<sup>2</sup>) |  |  |
| Protein (min/max/mean) | 3.58/162.64/85.75 | 11.57/163.90/89.45 |
| Nucleotide (min/max/mean) |  | 125.85/153.30/136.19 |
| Ligand (min/max/mean) | 156.12/168.30/162.21 | 130.67/156.31/143.49 |
| R.m.s deviations |  |  |
| Bond length (Å) | 0.003 | 0.005 |
| Bond angles (Å) | 0.642 | 0.762 |
| **Validation** |  |  |
| MolProbity score | 1.29 | 1.84 |
| Clash score | 1.74 | 5.16 |
| Ramachandran plot |  |  |
| Outliers (%) | 0.23 | 0.56 |
| Allowed (%) | 5.04 | 9.79 |
| Favored (%) | 94.73 | 89.65 |
| Rotamer outliers (%) | 0.52 | 1.02 |
| C-beta deviations (%) | 0.00 | 0.00 |

POT1(ESDL) is held in a single conformation stretched along the entire length of CST and buries a total of 6,623 Å^2^ of solvent-accessible surface area (Fig. 2D-E). The structures of the apo and ssDNA-bound complexes had identical overall interactions between POT1(ESDL) and CST, but the addition of ssDNA allowed resolution of POT1^OB-1^, which binds ssDNA in conjunction with POT1^OB-2^ (Fig. 2C, E). Because the region of the map containing the POT1 N-terminal OB-folds was at lower resolution (Fig. S3H; Fig. S4H), we could only dock in the existing crystal structure of POT1 bound to 5’-TTAGGGTTAG-3’ (PDB 1XJV ^48^) and could not assign a register to the DNA or observe additional nucleotides of the added [GGTTAG]_3_ oligonucleotide. Without ssDNA, POT1^OB-1^ appears to be flexibly attached and thus remains unresolved in the map of the apo complex (Fig. 2B; Fig. S3).

### Phosphorylation of POT1 mediates the interaction with CST

The hinge region of POT1(ESDL) that contains the ESDL insertion (aa 308-322) sits in a positively charged cleft of Ctc1^OB-D^ (Fig. 3A; Fig. S6A). Although there is connectivity in the cryo-EM map, we could not unambiguously resolve the side chain positions, suggesting this interaction is mediated by weaker interactions (Fig. 3A; Fig. S6A). However, this region of the POT1 hinge is negatively charged and contains multiple serine and aspartate residues that complement the basic cleft of Ctc1, and it is this charge complementarity that likely drives the interaction (Fig. 3A; Fig. S6A). Notably, the ESDL insertion does not make specific contacts with Ctc1, suggesting that the observed complex is not the result of an artificial mouse-human chimeric interaction. The abundance of Ser residues in this region suggests that this interaction may be regulated by phosphorylation of the POT1 hinge.

**Fig. 3.**
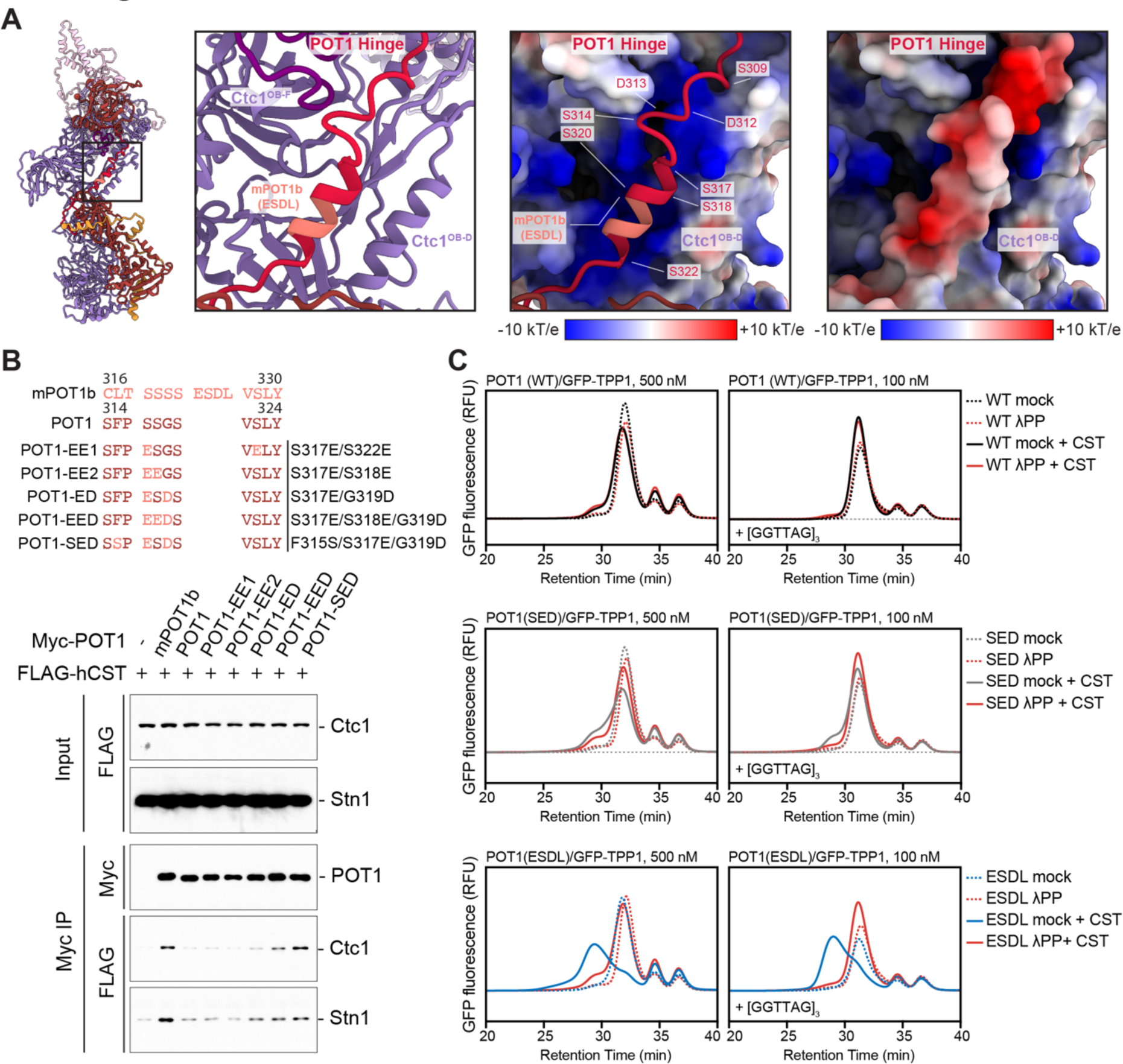
Phosphorylation-dependent interaction of POT1(ESDL) with CST. (**A**) Left panels: Structure of apo CST–POT1(ESDL)/TPP1 (left) and close-up view of the interaction of the POT1 hinge with Ctc1^OB-D^ (right). Right panels: Surface electrostatic potential analysis of the POT1(ESDL) hinge interacting with Ctc1^OB-D^. The POT1(ESDL) hinge is shown in cartoon (left) and surface (right) representation with electrostatic potential colored from red (−10 k_B_T/e) to white (0 k_B_T/e) to blue (+10 k_B_T/e) (see also Fig. S9). (**B**) Immunoblots of anti-Myc co-IPs of Myc-tagged POT1 constructs and FLAG-tagged CST from co-transfected 293T cells showing enhanced interaction with CST upon increasing negative charge of amino acids in the POT1 hinge. Immunoblots were probed with anti-Myc and anti-FLAG antibodies. The sequences of the altered POT1 alleles in comparison to POT1 and mPOT1b are displayed above the immunoblot. The first three lanes also appear in Fig. 1A. They are reproduced here as the controls for the experiment, which was performed simultaneously. (**C**) FSEC analysis of the phosphorylation-dependent interaction between CST and POT1(WT)/GFP-TPP1 (top, blue), POT1(SED)/GFP-TPP1 (middle, purple), and POT1(ESDL)/GFP-TPP1 (bottom, red) in the presence (right) or absence (left) of telomeric ssDNA (Fig. S9B). Traces without CST are shown as dashed lines, and traces containing POT1/TPP1 that have been treated with λ protein phosphatase (λPP) are shown in orange. RFU: relative fluorescence units. The chromatograms for the WT and ESDL mock samples also appear in Fig. 1F-G. They are reproduced here as the controls for the phosphatase experiment, which was performed simultaneously.

Indeed, negative charge substitutions in the hinge enhanced the POT1–CST interaction as measured by co-IP and with purified proteins *in vitro* (Fig. 3B-C). Of the charge substitution mutants, POT1-SED (F315S/S317E/G319D) showed the most robust interaction with CST in co-IP (Fig. 3B) but did not interact as well with CST as POT1(ESDL) *in vitro* (Fig. 3C). POT1-SED introduces a similar amount of negatively charged amino acids as POT1(ESDL), so the *in vitro* result suggests that the ESDL adds extra function in addition to simply introducing three charged residues.

To test the role of phosphorylation, we treated the POT1/TPP1 proteins with λ protein phosphatase. The CST–POT1(ESDL)/TPP1 interaction was severely diminished by dephosphorylation of POT1(ESDL)/TPP1, suggesting that the ESDL primarily functions to enhance the phosphorylation of human POT1 in both insect cells (Fig. 3C; Fig. S6B1) and 293T cells (Fig. 1C) rather than to directly interact with CST. The POT1-SED interaction is also diminished by phosphatase treatment, but to a lesser degree than in POT1(ESDL). Both POT1-SED and POT1(ESDL) return to a similar baseline interaction that could be attributed to the phosphorylation-independent negative charge added by each (Fig. 3C). The propensity of the mPOT1b hinge residues to be phosphorylated was also shown in a kinase assay in which HeLa cell extract was incubated with nitrocellulose membrane-bound synthetic peptides (Fig. S6C). The hinge peptides of human POT1 and mPOT1a showed less phosphorylation in the *in vitro* kinase assay than mPOT1b, consistent with their reduced ability to bind CST in co-IP experiments.

### POT1^OB-3^ forms the primary interface with Ctc1

Human POT1 directly interacts with Ctc1 at two sites separate from the ESDL insertion (Fig. 2D-E). The primary interface forms between POT1^OB-3^, Ctc1^ARODL^, and Ctc1^OB-D^ and buries 2,012 Å^2^ of solvent-accessible surface area (Fig. 4A-B; Fig. S6D). This interface is formed by the C-terminal region of the POT1 hinge (aa 322-341), which folds back on POT1^OB-3^ to create the surface to which Ctc1 binds. Part of the hinge has been previously crystallized ^47^, but in another crystal structure it was too flexible to be resolved ^46^ (Fig. 4A). In our structure, the hinge is pinned to POT1^OB-3^ at two sites (sites (i) and (ii)) of intramolecular interactions that limit its flexibility (Fig. 4A-B). POT1 Ser322 makes the furthest N-terminal self-interaction with POT1^OB-3^ at site (i), forming a salt bridge with Arg367 to create a linchpin that locks aa 322-341 in place (Fig. 4B; Extended Data Fig. 6D). Ser322 is mutated to Leu in CP ^27^, which would disrupt the electrostatic interaction and potentially disrupt motifs important for the phosphorylation of the POT1 hinge. This interaction is conserved, as mutation of the corresponding Ser328 in mPOT1b abrogates CST binding (Fig. S6E). At site (ii), aa 335-337 of the hinge interact with the C-terminus of POT1 (aa 627-629) in an anti-parallel β-strand configuration, as previously observed ^47^ (Fig. 4B; Fig. S6D). In the context of the full complex, these two points of contact secure the hinge against POT1^OB-3^ (Fig. 4A-B).

**Fig. 4.**
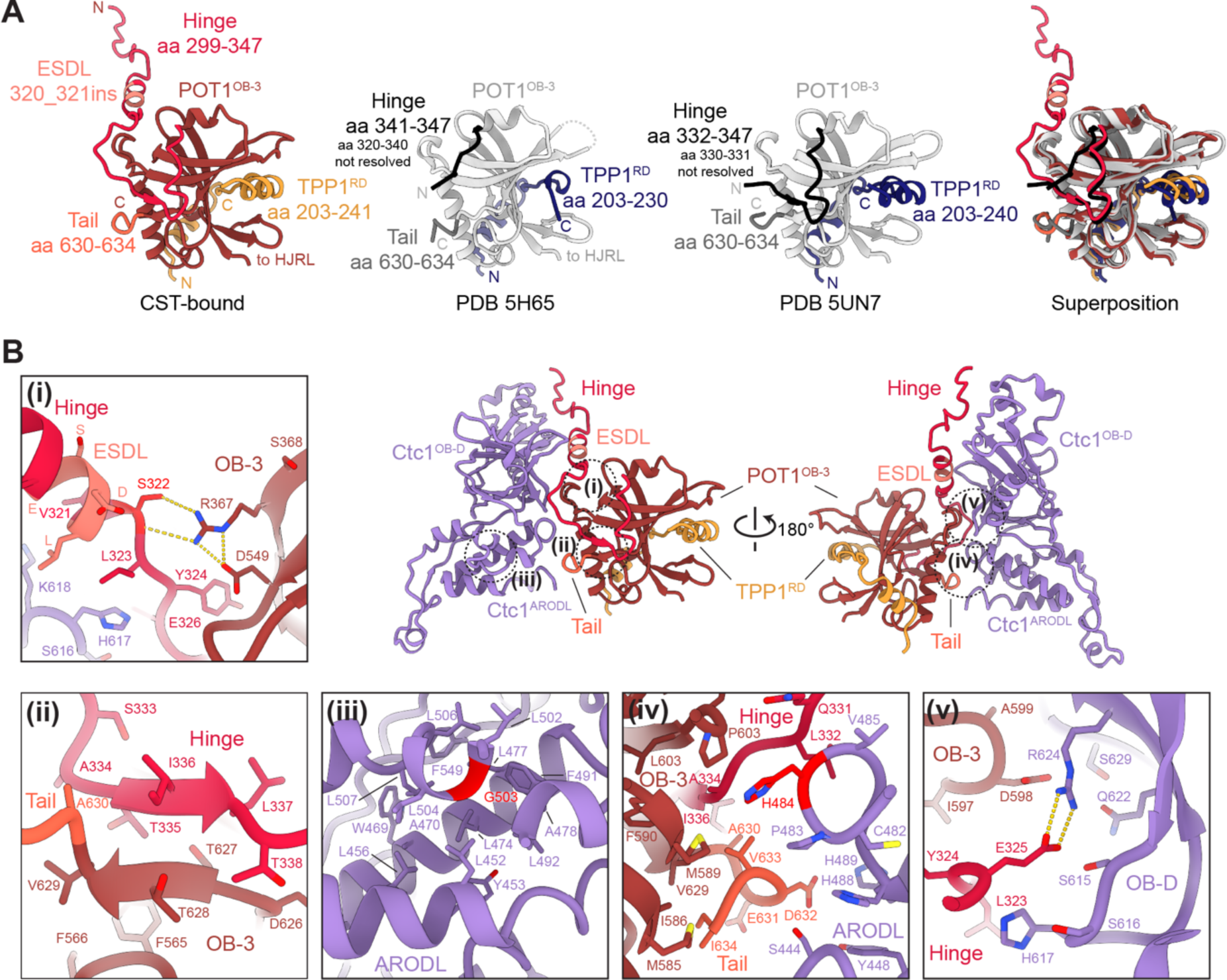
CP mutations map to the primary Ctc1–POT1 interface. (**A**) Comparison of CST-bound POT1^OB-3^/TPP1 (this study, colored) to unbound POT1^OB-3^/TPP1 (PDB 5H65/5UN7 ^46,47^, grayscale). (**B**) Close-up views of sites of interest at or near the interface between POT1 and Ctc1 (see also Fig. S6). Residues0 colored in bright red are mutated in CP. (i) Self-interaction between POT1 Ser322 and Arg367. Salt bridges are shown as yellow dashed lines. CP mutation S322L is predicted to disrupt this salt bridge and affect phosphorylation of the hinge. (ii) Self-interaction between POT1 hinge and C-terminus. (iii) Localization of Gly503 in the hydrophobic core of Ctc1^ARODL^ predicts a destabilizing effect of the G503R CP mutation. (iv) Primary hydrophobic interface between POT1 hinge, POT1^OB-3^, and Ctc1^ARODL^. CP mutation H484P is predicted to disrupt the hydrophobic stacking interactions with POT1 Pro603 and Ctc1 His488 and Pro483. (v) Salt bridges between POT1 hinge residue Glu325 and Ctc1^OB-D^ residue Arg624.

Our structure also provides insight into the Ctc1 G503R CP mutation ^20,49,50^. Although Gly503 is not at the Ctc1–POT1 interface, it is in the hydrophobic core of Ctc1^ARODL^ at site (iii). An arginine substitution would destabilize Ctc1^ARODL^, which forms the primary interaction with POT1 (Fig. 4B; Fig. S6D). This interaction is also conserved, as CST bearing the Ctc1 G503R mutation loses its interaction with mPOT1b (Fig. S6F). A loop in Ctc1^ARODL^, which interacts with POT1, is the location of another CP mutation, H484P ^28^. At site (iv), His484 participates in a network of hydrophobic stacking interactions with Ctc1 Pro483, Ctc1 His488, and POT1 Pro603. This interface is stabilized by hydrophobic packing of the Ctc1^ARODL^ loop against the POT1 tail (aa 630-634), POT1^OB-3^, and its hinge (Fig. 3B).

The end of the POT1 tail (632-DVI-634 in humans) was previously implicated in CST binding by mPOT1b ^30^. Substitution of the sequence in mPOT1b (DII) to the mPOT1a sequence (NVV) diminished the interaction with CST ^30^, and our structure explains the importance of the bulky hydrophobic residues at this site. The structure also shows that POT1 Asp632 is at an appropriate distance of Ctc1 His489 to form a salt bridge, which would further stabilize the complex (Fig. 4B; Fig. S6D). Finally, there is an additional salt bridge between POT1 Glu325 and Ctc1 Arg624 at site (v), which are in the POT1 hinge and Ctc1^OB-D^, respectively (Fig. 4B; Fig. S6D).

### POT1 blocks the CST ssDNA-binding site

The second OB fold of POT1, POT1^OB-2^, also interacts with CST. In contrast, POT1^OB-1^ does not directly interact with CST although its position could be determined in the ssDNA-bound complex (Fig. 2C, E). The resolution of the POT1^OB-2^–CST interface was in the 6-Å range (Fig. S3H; Fig. S4H), but still allowed for visualization of the docked structures. POT1^OB-2^ sits between Ctc1 and the C-terminal lobe of Stn1 (Stn1^C^) at Ctc1’s ssDNA-binding site ^16^ (Fig. 5A). The POT1 ssDNA-binding interface is facing outwards and is bound to DNA rather than facing inwards toward the Ctc1 ssDNA-binding interface (Fig. 2E; Fig. 5A). Helix αD of POT1 directly superposes with the DNA observed in the cryo-EM structure of CST ^16^ and the POT1 hinge runs through the DNA-binding site towards the basic cleft in Ctc1^OB-D^ (Fig. 5A; Fig. 3A). Stn1^C^, which can bind Ctc1 in multiple configurations, is in a position consistent with the structure of monomeric CST in the absence of ssDNA ^16^ (Fig. 5A). There was additional low-resolution density of a flexible loop in Ctc1 (aa 909-927), which has not been resolved in other structures of CST, and we used poly-alanine stubs to indicate the presence of this loop. The low resolution indicates that this loop contacts POT1^OB-2^ weakly (Fig. 5A).

**Fig. 5.**
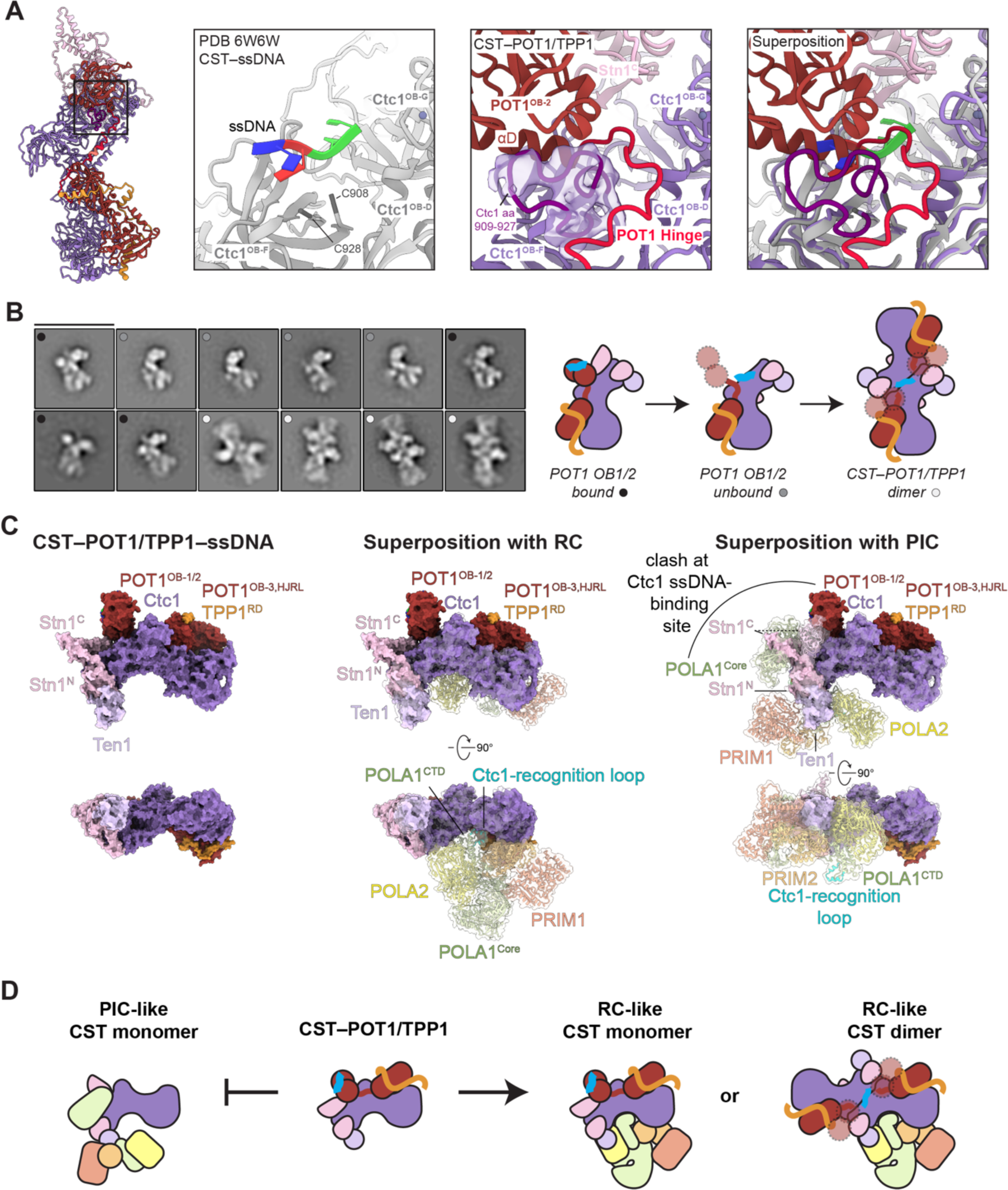
POT1^OB-2^ interacts with CST’s ssDNA-binding site, precluding formation of the CST–Polα/Primase PIC. (**A**) Interaction between POT1^OB-2^ and Ctc1 at the CST ssDNA-binding interface. (Left) The black box indicates the region of the apo CST–POT1(ESDL)/TPP1 model shown in the right panels. (Right) Zoom views of the interface. The first panel shows the ssDNA-bound CST structure (PDB 6W6W ^16^, gray with DNA colored). The second panel shows the same view of the CST–POT1(ESDL)/TPP1 structure (this study, colored). Ctc1 aa 909-927 are modeled as poly-alanine stubs, and no ssDNA is bound to Ctc1. The two structures are superposed in panel three. (**B**) Negative-stain EM 2D averages (left) of ssDNA-bound CST–POT1(ESDL)/TPP1 showing three major conformations depicted by the cartoon schematics (right). 2D averages are flipped or rotated by 90° increments from Fig. S4 into similar orientations for ease of comparison and are sorted by number of particles per class from the most populous class at top left to the least populous class at bottom right. The 2D averages that correspond to each cartoon state are indicated with black, gray, or white circles. The scale bar represents 333 Å (See also Fig. S4). (**C**) The CST–POT1(ESDL)/TPP1–ssDNA complex (left) and its superposition with the CST–Polα/Primase recruitment complex (RC, middle) and pre-initiation complex (PIC, right). The CST–POT1(ESDL)/TPP1–ssDNA complex is shown as an opaque surface, and CST–Polα/Primase complexes are shown in cartoon representation with transparent surfaces. Orthogonal views show that POT1(ESDL)/TPP1 binding does not obstruct the major interface of the RC, but POT1^OB-1/2^ and Stn1^C^ would interfere with binding of the POLA1 catalytic core to the CST ssDNA-binding site in the PIC (see also Fig. S9). (**D**) Cartoon schematics showing how CST–POT1/TPP1 is incompatible with PIC formation but could form RC-like complexes in both monomeric and dimeric forms.

The particles of the ssDNA-bound CST–POT1/TPP1 dataset were more heterogenous than those in the apo dataset; a subset of particles appeared to be missing density for the POT1 N-terminal OB-folds and Stn1^C^ (Fig. S2E; Fig. S4D; Fig. S5A). Loss of this density suggests that ssDNA is bound to Ctc1 rather than to POT1 as observed in the major conformation described above. Although POT1 has a greater affinity for telomeric ssDNA than CST ^51,52^, the reconstitution was performed with an excess of [GGTTAG]_3_ that could result in both proteins being bound to DNA. The POT1 C-terminus remains bound in these particles, observed in both negative-stain EM and cryo-EM (Fig. 5B; Fig. S2E; Fig. S4D; Fig. S5A). We could not obtain a high-resolution map for this conformation, likely because monomeric ssDNA-bound CST is flexible ^16^. It appears that the POT^OB-1/2^ and Stn1^C^-binding were important for stabilizing CST to allow high-resolution structure determination (Fig. 2B-C; Fig. S3-4) and loss of these interactions resulted in greater conformational heterogeneity (Fig. S5A). The negative-stain EM data suggest that this conformation is an intermediate in the dimerization of CST–POT1/TPP1 (Fig. 5B). CST is known to dimerize in the presence of ssDNA ^16^, and CST dimers clearly bound by POT1/TPP1 are observed in negative-stain EM 2D averages (Fig. 5B; Fig. S2E). CST dimerization requires displacement of POT1^OB-1/2^ and Stn1^C^ ^16^, but the POT1 C-terminus appears to be still attached by the POT1^OB-3^ and hinge interactions (Fig. 3-4). The CST–POT1/TPP1 dimers, which were a minor fraction of the particles observed in negative-stain EM averages (6%), could not be found in the cryo-EM dataset (Fig. S4), indicating that they may not withstand the vitrification process.

### POT1/TPP1 can recruit CST–Polα/Primase in the RC but not in the PIC state

The ssDNA-binding interface of CST is critical for formation of the CST–Polα/Primase PIC, in which the POLA1 catalytic core contacts the telomeric ssDNA template and is stabilized by Stn1^C^, which is in a different position relative to the monomeric and DNA-bound states of CST ^19^. POT1/TPP1 binding to CST is incompatible with Polα/Primase binding in a PIC-like conformation when POT1^OB-1^, POT1^OB-2^, and Stn1^C^ are engaged (Fig. 5C; Fig. S7A-B; Video S2). Therefore, we propose that POT1/TPP1 does not bind CST–Polα/Primase in the PIC conformation. In contrast, the major interface between Ctc1 and Polα/Primase in the auto-inhibited RC-like conformation is orthogonal to the POT1/TPP1 interface and is unobstructed, thus allowing for the formation of a POT1/TPP1-bound RC-like complex (Fig. 5C-D; Fig. S7A-B; Video S3). There is a minor clash between PRIM2 and POT1^HJRL^, but the cryo-EM structure of the RC was determined in cross-linking conditions, which limited the flexibility of the complex ^18^. This clash could be alleviated by the flexibility of CST relative to Polα/Primase observed in multi-body refinement analysis ^53^ of the CST–Polα/Primase RC complex (Fig. S7C; Videos S3-6). Structural heterogeneity of CST–Polα/Primase in an RC-like state was previously observed and proposed to be partially attributed to binding of the POLA2 N-terminal domain (POLA2^NTD^), which is attached by a flexible linker to POLA2 ^18^. Indeed, AlphaFold-Multimer predicts an interaction between the POLA2^NTD^ and Ctc1 (Fig. S7D-E), supporting the existence of flexible RC-like states observed in low-resolution cryo-EM maps ^18^.

What about the intermediate state in which the N-terminal OB folds of POT1 and Stn1^C^ are disengaged? We suggest that the OB-1/2-disengaged state represents a transient intermediate to the formation of a CST–POT1/TPP1 dimer (Fig. 5B). A CST–POT1/TPP1 dimer is likely to form at the telomere in the context of shelterin, which is fully dimeric and hence contains two copies of POT1/TPP1 ^37^. The high local concentrations of two CST molecules brought together by a shelterin dimer would favor dimerization. CST dimerization would also prevent PIC formation ^19^ and is compatible with the RC ^18^ (Fig. 5D).

## Discussion

The structures of CST bound to POT1/TPP1 reveal the mechanism by which human POT1 recruits and regulates CST (Fig. 6). This interaction is conserved between mice and humans and is likely conserved in other metazoan shelterin complexes, which all have POT1/TPP1 ^54^. Our data suggest that human POT1 toggles between the CST-recruitment function of mPOT1b and the telomere-protection function of mPOT1a using a switch in the phosphorylation state of residues in the POT1 hinge region. It will be important to determine which kinases and phosphatases are responsible for this switch and when and how phosphorylation is regulated.

**Fig. 6.**
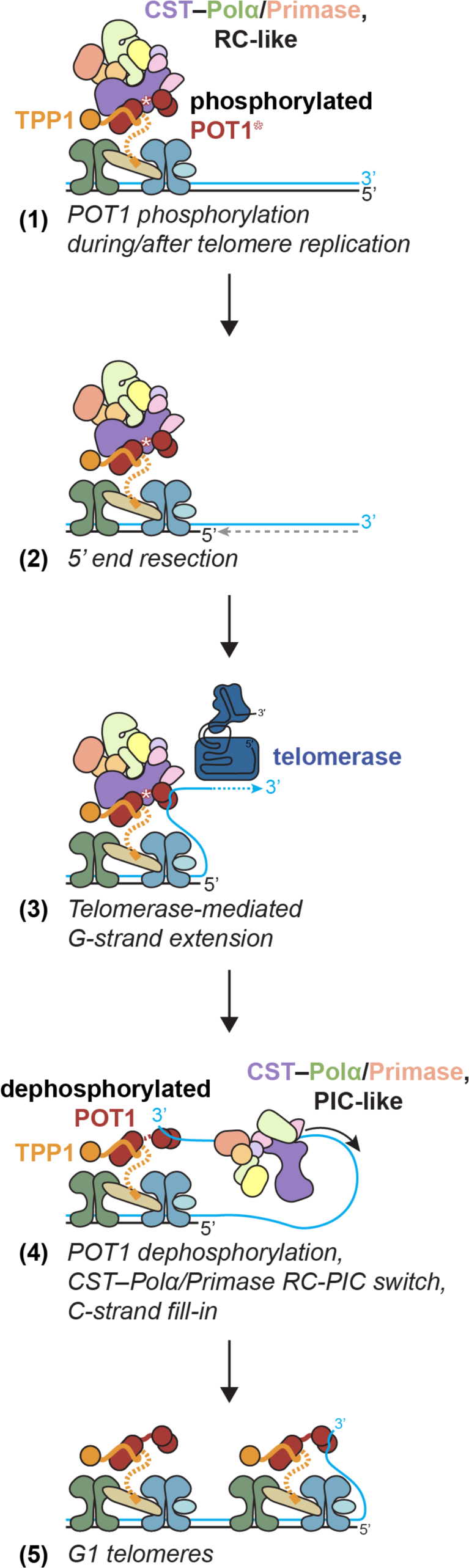
Model for telomere maintenance by shelterin and CST–Polα/Primase. Following DNA replication, CST–Polα/Primase is recruited to telomeres in the auto-inhibited RC-like state by phosphorylated POT1 (1). The POT1–CST interaction prevents activation of CST–Polα/Primase during 5’ end resection (2) and telomerase action (3). After telomerase has elongated telomeres, a cell cycle-dependent switch results in dephosphorylation of POT1 and subsequent release of CST–Polα/Primase to the telomeric ssDNA (4). When released, CST–Polα/Primase can readily form the PIC and execute fill-in synthesis, thereby completing telomere replication to form fully functional G1 telomeres (5).

Importantly, the results show how a subset of CP mutations affect telomeric CST recruitment, explaining the POT1 S322L genetic data and providing further evidence for the causative link between telomere maintenance defects and CP pathogenesis. Four CP mutations map to regions involved in CST–Polα/Primase recruitment: Ctc1 V665G maps to the primary interface observed in the RC, and POT1 S322L, Ctc1 H484P, and Ctc1 G503R affect CST–POT1 complex formation. Furthermore, CP mutations in Ctc1^OB-B^ are numerous and potentially affect the predicted interaction between POLA2^NTD^ and Ctc1, though further experiments are needed to confirm the AlphaFold-Multimer prediction. Together, these CP mutations highlight CST–Polα/Primase recruitment as a critical step in normal telomere maintenance. Once the kinase(s) responsible for regulation of this step is identified, it is possible that additional patient mutations will be discovered within the kinase or its regulatory factors.

The incompatibility of POT1/TPP1 binding with CST–Polα/Primase PIC formation corroborates the model in which CST–Polα/Primase is recruited by phosphorylated POT1 to the telomere in an auto-inhibited RC-like state ^18^. Such a regulatory role for POT1/TPP1 is consistent with the temporal delay of fill-in, which takes place after telomere replication, resection, and extension by telomerase (^31,55,56^; reviewed in (Cai and de Lange, in revision)). We suggest that dephosphorylation of POT1 is a critical step for the release of CST, formation of the PIC, and initiation of fill-in (Fig. 6).

We propose that the regulated activation of CST–Polα/Primase is needed to ensure that CST does not inhibit telomerase during the initial G-strand extension step (Fig. 6). Telomere length homeostasis is achieved by cis-acting inhibition of telomerase through a mechanism involving the shelterin proteins TRF1, TIN2, and POT1 ^8–10^. In addition, CST has emerged as a powerful inhibitor of telomerase both *in vivo* and *in vitro* ^36,49,57^ and it seems likely that CST acts downstream of TRF1/TIN2/POT1 although this has not been proven. The inhibitory activity of CST may have to be blocked during the telomere extension step when telomerase restores the 3’ overhang at leading-end telomeres. In this regard, it is of interest that the ssDNA-binding pocket of CST is blocked by POT1. Since the ssDNA binding of CST is required for telomerase inhibition *in vitro* ^57,58^, our data predict that CST is not capable of inhibiting telomerase while it is bound to POT1. The POT1 dephosphorylation step that allows CST–Polα/Primase to become activated may therefore also be required to allow CST to inhibit telomerase. Our data showing that CST does not interact with the N-terminal OB fold of TPP1 argue against an alternative model for telomerase regulation, in which CST competes with telomerase for binding to TPP1.

These findings open a new avenue for research on the regulation of telomere maintenance. Whereas telomere elongation by telomerase has been studied for more than 30 years, examination the equally important synthesis of the telomeric C-strand has lagged (pun intended). We expect that the identification of the kinases/phosphatases that control CST recruitment and its subsequent release will reveal how the cell cycle controls the critical last step in telomere maintenance.

## Acknowledgements

We thank J. Zinder for generating and optimizing the expression strategy of the wild-type POT1/TPP1 expression construct and for providing the λ protein phosphatase. T. Bakker is thanked for designing the TPP1-Stn1 fusion gene under the supervision of J. Zinder. We thank M. Ebrahim, J. Sotiris and H. Ng at the Evelyn Gruss Lipper Cryo-EM Resource Center of The Rockefeller University for assistance with cryo-EM data collection. Some grids were also frozen at the Electron Microscopy Resource Center of The Rockefeller University. SWC is supported by the David Rockefeller Graduate Program and an NSF Graduate Research Fellowship (1946429). HT is supported by an R50 grant from the NIH (5 R50 CA243771). This work is supported by grants to TdL from the NIH (5 R35 CA210036) and the Breast Cancer Research Foundation (BCRF-19-036).

## Author contributions

All authors contributed to the conception of the study and the writing of the manuscript. SWC performed and analyzed the experiments other than the co-IP and peptide phosphorylation assays, which were performed and analyzed by HT. SWC prepared figures. TW and TdL supervised all experiments and analysis.

## Competing interests

Titia de Lange is a member of the SAB of Calico Life Sciences LLC, San Francisco. The other authors have no conflicts to declare.

## Additional Information

Supplementary Information is available for this paper. Correspondence and requests for materials should be addressed to Titia de Lange.

## Methods

### DNA construct generation

The DNA constructs used in this study are listed in Table S1. For recombinant insect cell expression, the biGBac ^59^ vector system was used. All protein sequences are numbered according to the canonical sequence entry in UniProt. Recombinant bacmids were generated from the plasmids in Table S1 using MAX Efficiency DH10Bac competent cells (Gibco; Cat. 10361012) and transfected into Sf9 insect cells (Gibco; Cat. 11496015) with Cellfectin II Reagent (Gibco) to generate a P1 baculovirus stock. P1 baculovirus was amplified in adherent Sf9 insect cells to generate P2 and P3 stocks, and the P3 virus was used to infect Tni suspension insect cell culture (Expression Systems; Cat. 94-002S) for protein expression.

### Co-immunoprecipitation

293T cells were grown in DMEM with 10% (v/v) bovine calf serum (Hyclone), 2 mM L-glutamine (Gibco), 100 U/mL penicillin (Gibco), 0.1 mg/mL streptomycin (Gibco), 0.1 mM nonessential amino acids (Gibco). 4-5×10^6^ 293T cells were plated in a 10-cm dish 20-24 hours prior to transfection by calcium-phosphate precipitation. Transfections were done using plasmid DNA as indicated such that total DNA did not exceed 20 μg per plate. 48 hours later, 4×10^6^ cells were replated into a 15-cm dish and cultured for an additional 12-15 hours. The cells were harvested by trypsinization, rinsed with 1x phosphate-buffered saline (PBS; Corning), frozen once with liquid nitrogen (LN_2_), and resuspended in 0.5 mL lysis buffer (50 mM Tris-HCl pH 7.6, 150 mM NaCl, 1 mM EDTA, 10% (v/v) glycerol, 8 mM 2-mercaptoethanol (β-ME), 0.1-0.5% (v/v) NP-40, complete EDTA-free protease inhibitor cocktail (Roche), and PhosSTOP phosphatase inhibitor mix (Roche)), and incubated on ice for 1 hour. After centrifugation at 16,000×g for 10 min at 4°C, 1.5 μL of anti-Myc-tag 9B11 antibody (Cell Signaling, 2267), anti-Strep-tag antibody (Qiagen, 34850), or anti-HA-tag antibody (Cell Signaling, 3724) was added to the supernatant. Samples were nutated at 4°C for 4 hours. 20 μL of Protein G magnetic beads slurry (Cell Signaling, 9006), preincubated with 5% Bovine Serine Albumin (BSA) in PBS, was added and the sample was nutated at 4°C for an additional hour. Beads were washed 4 times at 4°C with lysis buffer containing 0.1% NP-40 and immunoprecipitated protein was eluted with 50 μL of 2x Laemmli buffer. Samples were boiled for 5 min before separation on SDS-PAGE.

### Peptide phosphorylation assay

The phosphorylation assay was done as previously described ^60^ with some modifications. 15-mer acetylated peptide arrays of the POT1 protein hinge regions (POT1 aa 313-327; mPOT1b aa 319-333; mPOT1a aa 319-333) were produced by Kinexus Bioinformatics Co. HeLa cells were grown in the same medium used for the 293T cell culture and were used for nuclear extract preparation. 6×10^7^ cells were harvested by trypsinization, rinsed with PBS, and washed two times with 2 mL of cytoplasm buffer (10 mM Tris-HCl pH 7, 10 mM NaCl, 3 mM MgCl_2_, 30 mM sucrose, and 0.5% NP-40). After centrifugation, the cell suspension was centrifuged, and the pellet was rinsed twice with 2 mL of CaCl_2_ buffer (10 mM Tris-HCl pH 7, 10 mM NaCl, 3 mM MgCl_2_, 30 mM sucrose, and 100 μM CaCl_2_). The pellet was resuspended in 1 mL of buffer C (20 mM Tris-HCl pH 7.9, 20% glycerol, 100 mM KCl, and 0.2 mM EDTA), and centrifuged at 20,000×g for 30 min at 4°C. The supernatant was collected and used as nuclear extract for the kinase assay. The nitrocellulose membrane of the peptide array was rinsed once with reaction buffer (20 mM Tris-HCl pH 7.9, 50 mM KCl, 10 mM MgCl_2_, 1 mM DTT, complete EDTA-free protease inhibitor cocktail, PhosSTOP), and was subsequently incubated in the 4 mL of reaction buffer containing 50 μM ATP, 50 μCi/μmol [r^32^P] ATP, and nuclear extract (16 μg protein) for 45 min at 30°C. EDTA (15 mM final concentration) was added to stop the reaction and the membrane was rinsed 10 times with cold 1 M NaCl. The membrane was rinsed once with wash buffer (4 M guanidine, 1% (v/v) SDS, and 0.5% β-ME) and exposed for radiography (Typhoon Biomolecular Imager, GE Healthcare).

### Protein expression and purification

50 mL of P3 baculovirus was used per 500 mL of Tni culture, infected at a cell density of 2×10^6^ cells/mL. The infected cells were grown in spinner flasks at 150 rpm for 72 hours at 27°C. Cells were harvested by centrifugation (500×g) and transferred to a syringe before flash freezing droplets in LN_2_. CST was purified as previously described ^18^. CST with the His_6_-MBP tag cleaved was only used in the negative stain analysis of CST– POT1(ESDL)/TPP1–ssDNA complex (Fig. S2E). After facing issues with aggregation and poor vitrification, the His_6_-MBP was retained in all other experiments to aid with protein stability.

For the POT1/TPP1 proteins, frozen pellets were lysed by cryogenic milling (Retsch) and the cryo-milled powder was resuspended in a buffer containing 50 mM HEPES pH 7.5, 350 mM NaCl, 5 mM β-ME, 5% glycerol, 0.05% (v/v) Tween-20, and 1 mM phenylmethylsulfonyl fluoride (PMSF), supplemented with complete EDTA-free protease inhibitor cocktail (Roche). The lysate was cleared by centrifugation at 4°C and 40,000×g for 1 h. Supernatants were incubated with end-over-end rotation for 1 hour at 4°C with Strep-Tactin Superflow high-capacity resin (IBA Lifesciences) equilibrated with gel-filtration buffer containing 20 mM HEPES pH 7.5, 300 mM NaCl, 0.1 mM tris(2-carboxyethyl)phosphine (TCEP), 0.05% Tween-20, and 5% glycerol and subsequently washed with 20-50 column volumes (CV) of the same buffer. Bound protein was eluted in the same buffer supplemented with 10 mM d-desthiobiotin, concentrated, and loaded onto a HiLoad Superdex 200 16/600 PG column (Cytiva) equilibrated with the same gel-filtration buffer.

The fusion CST–POT1/TPP1 complex was purified similarly, but after elution from the Strep-Tactin resin, it was concentrated to 500 μL and loaded on top of an 11-mL linear 10-30% glycerol gradient. Ultracentrifugation was carried out at 41,000 rpm in an SW 41 Ti rotor for 18 hours at 4°C. 500 μL fractions were manually collected and protein-containing fractions were identified by SDS-PAGE.

Protein-containing fractions were concentrated, flash frozen in LN_2_, and stored in aliquots at −80°C. Protein concentrations were measured on a NanoDrop-1000 spectrophotometer.

### Fluorescent size-exclusion chromatography analysis

Protein–protein interaction experiments were performed on a Superose 6 Increase gel-filtration column (Cytiva) equilibrated in buffer containing 20 mM HEPES pH 7.5, 150 mM NaCl, and 0.1 mM TCEP. Eluate from the column was passed through an RF-20A fluorescence detector (Shimadzu) operated with excitation and emission wavelengths of 280 nm and 340 nm, respectively.

Proteins were mixed to a final volume of 120 μL and incubated on ice for at least 30 minutes prior to injection of 100 μL on the column. The GFP-tagged POT1/TPP1 constructs were added at the specified concentrations, and [GGTTAG]_3_ ssDNA (when present) and CST were added at a twofold molar excess. Complex formation was evaluated by comparing the mobility on the gel-filtration column of POT1/TPP1 alone *versus* when mixed with CST.

For the dephosphorylation experiments, 10 μM POT1/TPP1 was incubated with 10 μM λ protein phosphatase (λPP, expressed and purified in-house) and 1 mM MnCl_2_ in the FSEC gel-filtration buffer. For mock treated samples, the λPP was omitted and replaced with buffer. Samples were incubated 8-16 hours. An aliquot for SDS-PAGE analysis was taken from each sample before they were diluted to prepare the FSEC experiments described above.

### Reconstitution of the ssDNA–CST–POT1/TPP1 complex

Purified His_6_-MBP-Ctc1/Stn1/Ten1 and His_6_-POT1(ESDL)/TwinStrep-GFP-TPP1 were mixed with [GGTTAG]_3_ ssDNA in 150 μL at final concentrations of 6 μM, 6 μM, and 9 μM, respectively, in a buffer containing 20 mM HEPES pH 7.5, 0.5 mM TCEP pH 7.5, and 1% glycerol. Based on the volume of the input proteins (which were purified at 300 mM NaCl), the NaCl was adjusted to a final concentration of 150 mM. The protein components were mixed first and incubated on ice for 1 hour prior to addition of the ssDNA. The protein and ssDNA were incubated on ice for 2 hours prior to loading on top of an 11-mL linear 10-30% glycerol gradient. Ultracentrifugation was carried out at 41,000 rpm in an SW 41 Ti rotor for 18 hours at 4°C. 500 μL fractions were manually collected and protein- and DNA-containing fractions were identified by SDS-PAGE and native PAGE.

### Negative-stain EM sample preparation, data collection, and image processing

Protein samples for negative-stain EM (3.5 μL drops, in a concentration range of 0.01-0.05 mg/mL) were adsorbed to glow-discharged carbon-coated copper grids with a collodion film, washed with three drops of deionized water, and stained with two drops of freshly prepared 0.7% (w/v) uranyl formate. Samples were imaged at room temperature using a Phillips CM10 electron microscope equipped with a tungsten filament and operated at an acceleration voltage of 80 kV. The magnification used for the CST-only samples corresponds to a calibrated pixel size of 3.5 Å. The magnification used for the CST–POT1(ESDL)/TPP1 samples corresponds to a calibrated pixel size of 2.6 Å. Particles were auto-picked using the Swarm picker (CST-only) or Gauss picker (CST– POT1(ESDL)/TPP1) in EMAN2 ^61^. Particle extraction and 2D classification were performed in RELION-3.1 ^62^.

### Cryo-EM sample preparation and data collection

Apo CST–POT1(ESDL)/TPP1 complex was frozen at a concentration of 0.075 mg/mL, corresponding to 0.2 μM of GFP-tagged protein, in buffer containing 20 mM HEPES pH 7.5, 150 mM NaCl, and 0.1 mM TCEP. The CST–POT1(ESDL)/TPP1–ssDNA complex was prone to aggregation following vitrification at the concentration used for the apo complex and faced challenges with inconsistent ice distribution across the grid. Thus, the CST–POT1(ESDL)/TPP1–ssDNA complex was diluted to 0.02 mg/mL, or 0.05 μM of GFP-tagged protein. The addition of 0.75 mM fluorinated Fos-choline-8 (Anatrace) resulted in even ice distribution.

3.5 μL of the samples were applied to Quantifoil R1.2/1.3 mesh Cu400 holey carbon grids covered with a graphene oxide support layer (EMS) that were glow-discharged for 5 s at 40 mA in an EMS100X glow discharge unit (EMS) and then blotted for 0.5-1 s (Blot Force −2; Wait Time 20 s; Drain Time 0 s) and plunge frozen in liquid ethane using a Vitrobot Mark IV (Thermo Fisher Scientific) operated at 4°C and 100% humidity. Cryo-EM imaging was performed in the Evelyn Gruss-Lipper Cryo-EM Resource Center at The Rockefeller University using SerialEM ^63^. Data collection parameters are summarized in Table 1.

For both complexes, data were collected on a 300-kV Titan Krios electron microscope equipped with a Cs corrector at a nominal magnification of ×105,000, corresponding to a calibrated pixel size of 0.86 Å (micrograph dimensions of 5,760×4,092 px) on the specimen level. Images were collected using a slit width of 20 eV on the GIF Quantum energy filter (Gatan) and a defocus range from −1 to −2.5 μm with a K3 direct electron detector (Gatan) in super-resolution counting mode. Three movie stacks were recorded per hole in a 3×3 matrix of holes acquired using beam-image shift with a maximum shift of 3.5 μm. Exposures of 1.2 s were dose-fractionated into 40 frames (30 ms per frame) with an exposure rate of 31 electrons/pixel/s (approximately 1.25 electrons per Å^2^ per frame), resulting in a total electron exposure of 50.3 electrons per Å^2^.

### Cryo-EM data processing

For all datasets, movie stacks were motion-corrected with the RELION-3.1 ^62^ implementation of MotionCor2, and motion-corrected micrographs were manually inspected and curated (graphene oxide coverage of grids was inconsistent) prior to CTF parameter estimation with CTFFIND-4 ^64^ implemented in RELION-3.1.

To generate templates and an initial model for the apo complex, particles were automatically picked without a template using Gautomatch and extracted (352 px box, binned 4-fold to 88 px) in RELION-3.1. After 2D classification and cleanup to remove contaminating ice particles, the cleaned particle stack was imported into cryoSPARC (v4.1.1) and 2D classification was performed. The best 2D classes (containing ∼150,000 particles) were used to generate an initial model that was used as a 3D reference. The motion-corrected micrographs were then imported into cryoSPARC, and CTF parameter estimation was performed with CTFFIND-4 ^64^ implemented in cryoSPARC. Particles were automatically picked using the Template Picker with the initial model as a 3D reference. Particles were extracted and Fourier-cropped to speed up image processing (352 px box → 256 px box) and processed using the described image processing pipeline (Fig. S3). Three rounds of heterogeneous refinement were performed with one good reference and three noise “decoy” references to filter out junk particles. After the number of particles in the junk classes dropped to 15% or below, the good particle stack was further classified using heterogeneous refinement into four classes using a single reference. The 372,930 particles in the best class were re-extracted at full-size (352 px box). There was additional heterogeneity, particularly in the region of Stn1^C^ and POT1^OB-1/2^, so further unsupervised heterogeneous refinement was used to classify particles with fully intact Stn1^C^ and POT1^OB-1/2^. The final stack of 132,356 particles was refined and sharpened to a nominal resolution of 3.6 Å using the Non-uniform Refinement job in cryoSPARC and local resolution of the map was estimated in cryoSPARC. The reported 3.9-Å resolution was calculated with Phenix (phenix.validation_cryoem) using the gold-standard Fourier shell correlation (FSC) between half-maps (Fig. S3I), and FSC plots and sphericity values ^65^ of the reconstruction were calculated using the 3D-FSC server (https://3dfsc.salk.edu/).

For the ssDNA-bound complex, the motion-corrected micrographs were imported into cryoSPARC, and CTF parameter estimation was performed with the Patch CTF Estimation job. The apo complex map was used as a 3D template for particle picking with the Template Picker job. Particles were extracted and Fourier-cropped to speed up image processing (352 px box → 128 px box) and processed using the described image processing pipeline (Fig. S4). The 2D-class averages of the extracted particles and negative-stain EM analysis of the complex (Fig. 4) revealed heterogeneity. This heterogeneity could be divided into three categories of particles: fully intact CST– POT1/TPP1 complexes that resembled the apo complex and contained bound Stn1^C^ and POT1^OB-1/2^, half complexes in which Stn1^C^ and POT1^OB-1/2^ were disengaged but the POT1 C-terminus was bound, and particles containing CST without POT1/TPP1. Thus, three 3D references were generated from the apo complex by segmenting the map in UCSF Chimera. The references are shown in Fig. S4. Similar to the apo complex, two noise decoy references were used in addition to the three real references to perform iterative heterogeneous refinement until the noise classes contained fewer than 10% of the particles. An additional heterogeneous refinement with the three different references was performed prior to re-extracting the particles with Stn1^C^ and POT1^OB-1/2^ bound at full size. There was still some heterogeneity in that region, so the map was segmented to generate a mask around the Ctc1 C-terminus, Stn1, Ten1, and POT1^OB-1/2^. This mask was used to perform focused 3D classification into four classes without alignment in cryoSPARC. Two classes contained density for both Stn1^C^ and POT1^OB-1/2^ and were pooled into a final stack of 76,359 particles. The other two classes were missing density for one of the components. The final particle stack was refined and sharpened to a nominal resolution of 4.0 Å using the Non-uniform Refinement job in cryoSPARC and local resolution of the map was estimated in cryoSPARC. The reported 4.3-Å resolution was calculated with Phenix (phenix.validation_cryoem) using the gold-standard FSC between half-maps (Fig. S4I), and FSC plots and sphericity values ^65^ of the reconstruction were calculated using the 3D-FSC server (https://3dfsc.salk.edu/).

We also attempted to continue processing the complex in which ssDNA was presumably bound to CST (Fig. S5A). In this complex, only the C-terminus of POT1 is bound to CST. The classes generated by this reference complex were pooled at two steps and combined into a stack containing 502,366 particles. These particles were first aligned using the homogeneous refinement job in cryoSPARC. This map contained an additional cylindrical density seen at low thresholds that was attributed to TPP1^OB^ (Fig. S5B) but this density disappeared with heterogeneous refinement. Focused 3D classification revealed that there were few particles truly missing the Stn1^C^ and POT1^OB-1/2^ as cartooned in Fig. 5B, but the association was weak. Neither heterogeneous refinement nor focused 3D classification without alignment were able to further classify the particles or align to higher-resolution features. This is likely due to the intrinsic flexibility of CST, for which a high-resolution structure of a monomeric complex without other interacting factors could not be determined ^16^. It appears that when bound, the Stn1^C^ and POT1^OB-1/2^ interactions stabilized CST to allow for high-resolution structure determination.

### Model building and refinement

The map of the apo complex was used for atomic model building, as it had a higher overall resolution and better quality. The AlphaFold2 ^44,45^ database model of Ctc1 (AF-Q2NJK3) was merged with the cryo-EM structure of CST (PDB 6W6W ^16^) as the starting model for CST. The X-ray crystal structures of POT1^OB-3/HJRL^/TPP1^RD^ (PDB 5H65 ^46^) and the POT1 OB-folds bound to ssDNA (PDB 1XJV ^48^) were used as starting models for the POT1/TPP1 components. For the apo complex, CST, POT1^OB-2^, and POT1^OB-3/HJRL^/TPP1^RD^ were first rigid-body docked into the map using UCSF Chimera. The model was then manually inspected in Coot ^66^, where additional density of the hinge region of POT1 was observed. There was clear density for aa 322-347 that allowed unambiguous manual modeling of the hinge self-interactions with POT1^OB-3^. The N-terminal region of the hinge (aa 299-321 and the ESDL insertion) showed connected Cα backbone density, but many side-chain placements were ambiguous. Because this region of the hinge makes a weak interaction that appears to be mediated primarily by charge complementarity rather than specific interactions, the exact side-chain positions were not critical to the analysis. Thus, we first modeled the hinge Cα backbone into the density and observed that it spanned the distance between the preceding and following regions of the chain. Then, we assigned the sequence in register with the rest of the protein. The key reason for modeling the side-chains in residues 308-321 was to show the abundance of serine and aspartate residues in this region. Residues 301-307 were modeled as poly-alanine stubs, as this part of the chain runs in a groove of Ctc1 and we did not want to over- or mis-interpret specific interactions. The resolution of POT1^OB-2^ was not high, so POT1^OB-1^ was only flexibly fitted into the density using ISOLDE ^67^ after rigid-body docking. For Ctc1, the AlphaFold2 model fit well overall into the density and required only few manual adjustments. One region of Ctc1 appeared to contact POT1 weakly (aa 909-927). This region of Ctc1 has not been resolved in previous experimental structures and AlphaFold2 predicts this loop with low confidence. There was clear density near the predicted position of the loop, so it was approximated with poly-alanine stubs. The map regions containing Stn1 and Ten1 were also at lower resolution, so they were docked in and fitted similar to POT1^OB-2^. Iterative real-space refinement in Phenix (phenix.real_space_refine) and manual model adjustment in ISOLDE ^67^ was used to correct the most egregious geometry errors, though some errors could not be fixed in a way supported by the experimental map. The model and maps were validated in Phenix (phenix.validation_cryoem) ^68^.

For the ssDNA-bound complex, the apo complex model was used as the starting model. POT1^OB-2^ from the apo complex was replaced with a rigid-body docking of POT1^OB-1/2^ and a 5’-TTAGGGTTAG-3’ DNA from the X-ray crystal structure (PDB 1XJV ^48^). Because the overall resolution of the map of the ssDNA-bound complex was lower, but the overall interaction and map density was similar between the apo and ssDNA-bound complex, we treated the apo model as relatively correct. The apo model was fitted to the ssDNA-bound complex map using iterative real-space refinement in Phenix (phenix.real_space_refine) and some manual model adjustment in ISOLDE ^67^. The resolution of POT1^OB-1/2^ was low, so it was modeled by flexible fitting of the crystal structure and some attempts were made to correct its geometry in ISOLDE ^67^. ssDNA was clearly present, but the resolution was too low to assign the register or see if more than 10 nt were bound relative to the crystal structure. Thus, only the 5’-TTAGGGTTAG-3’ from the crystal structure was retained in the final model. The model and maps were validated in Phenix (phenix.validation_cryoem) ^68^.

For PDB deposition, the POT1 ESDL insertion was notated using insertion code as an insertion after residue 320 (E – 320A; S – 320B; D – 320C; L – 320D) to maintain the canonical numbering of the human POT1 sequence. The TPP1–Stn1 fusion in the apo complex required deposition as a single chain, so the numbering corresponds to the fusion. However, individual mappings to TPP1 and Stn1 within the chain are annotated in the deposition.

### Multi-body refinement analysis of CST–Polα/Primase RC flexibility

The dataset used for RC analysis was the same as previously published ^18^ (EMPIAR-11131) and reprocessed in RELION-3.1 with the same particle coordinates and particles extracted. The final stack used for multi-body refinement contained 203,045 particles (300 px box; 1.08 Å/px). Masks for Polα/Primase (body 1) and CST (body 2) were generated by segmentation of the RC map in UCSF Chimera and mask creation in RELION-3.1. Multi-body refinement was performed with default parameters ^53^. Angular/translational priors of 10°/2px and 20°/5px were used for Polá/Primase and CST, respectively. Volume series movies were generated in UCSF ChimeraX ^69^.

### AlphaFold-Multimer complex structure prediction

AlphaFold-Multimer (v1.0) was used on the COSMIC2 server ^70^ with default settings (full_dbs). For the Ctc1^OB-A/B/C^–TPP1^TID^ complex, the input sequence file contained Ctc1 aa 1-410 and TPP1 aa 361-458. For the Ctc1–POLA2 complex, the input sequence file contained Ctc1 aa 1-1217 and POLA2 aa 1-158. Flexible regions with low predicted confidence were hidden from the figures.

### Structure analysis and visualization

The apo CST–POT1(ESDL)/TPP1 complex was used for all interaction analysis, as the model and map quality was superior to that of the ssDNA-bound complex. The additional POT1^OB-1^ and telomeric DNA in the ssDNA-bound complex did not interact with CST. UCSF Chimera ^71^, UCSF ChimeraX ^69^, and PyMOL (Schrödinger) were used to analyze the maps and models. Figures were prepared using UCSF ChimeraX and Adobe Illustrator.

## Data availability

The cryo-EM maps have been deposited at the Electron Microscopy Data Bank under accession codes EMD-40659 (apo CST–POT1/TPP1 complex) and EMD-40660 (ssDNA–CST–POT1/TPP1 complex), and the coordinates have been deposited in the Protein Data Bank under accession codes PDB 8SOJ and PDB 8SOK, respectively.

---

**Supplemental Fig. S1.**
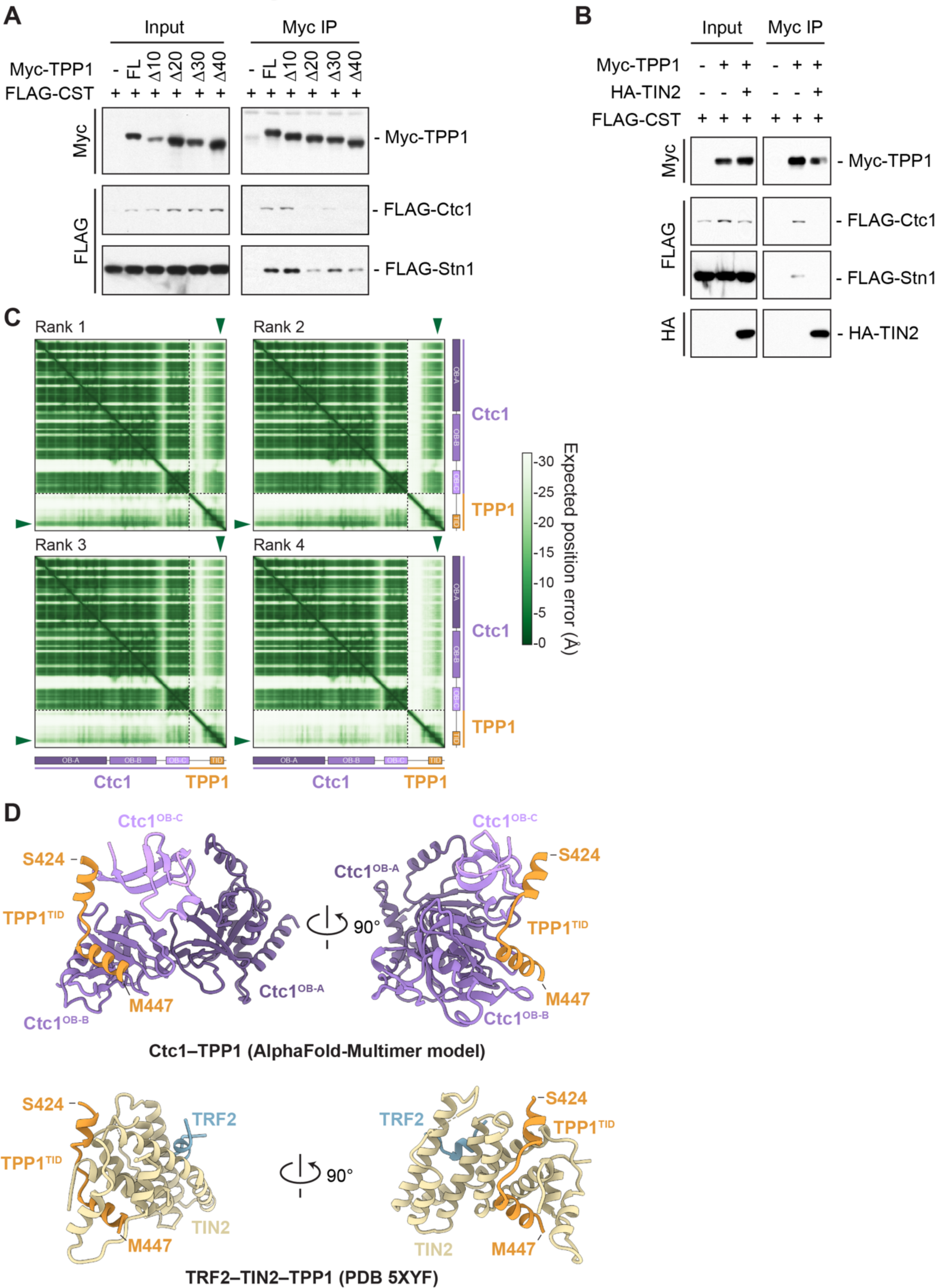
Analysis of the CST–TPP1 interaction. (**A**) Immunoblots of anti-Myc co-IPs of Myc-tagged TPP1 constructs with the indicated C-terminal truncations and FLAG-tagged CST from co-transfected 293T cells showing that the C-terminus of TPP1 is required for the CST–TPP1 interaction. Immunoblots were probed with anti-Myc and anti-FLAG antibodies. (**B**) Immunoblots on anti-Myc co-IPs of Myc-tagged TPP1, HA-tagged TIN2, and FLAG-tagged CST from co-transfected 293T cells showing that TIN2 competes with CST for TPP1 interaction. Immunoblots were probed with anti-Myc, anti-FLAG, and anti-HA antibodies. (**C**) Predicted aligned error (PAE) plots of the top four (of 5) ranked AlphaFold-Multimer ^40^ models for Ctc1–TPP1. Green arrowheads indicate high confidence in the position prediction of TPP1 relative to Ctc1. All 5 top-ranked models predicted the interaction with high confidence. (**D**) AlphaFold-Multimer model of Ctc1 OB-folds A, B, and C bound to TPP1^TID^ compared to the crystal structure of TRF2–TIN2–TPP1^TID^ (PDB 5XYF ^39^). TPP1^TID^ is predicted to bind to Ctc1 and TIN2 using the same peptide, providing a structural basis for their competition.

**Supplemental Fig. S2.**
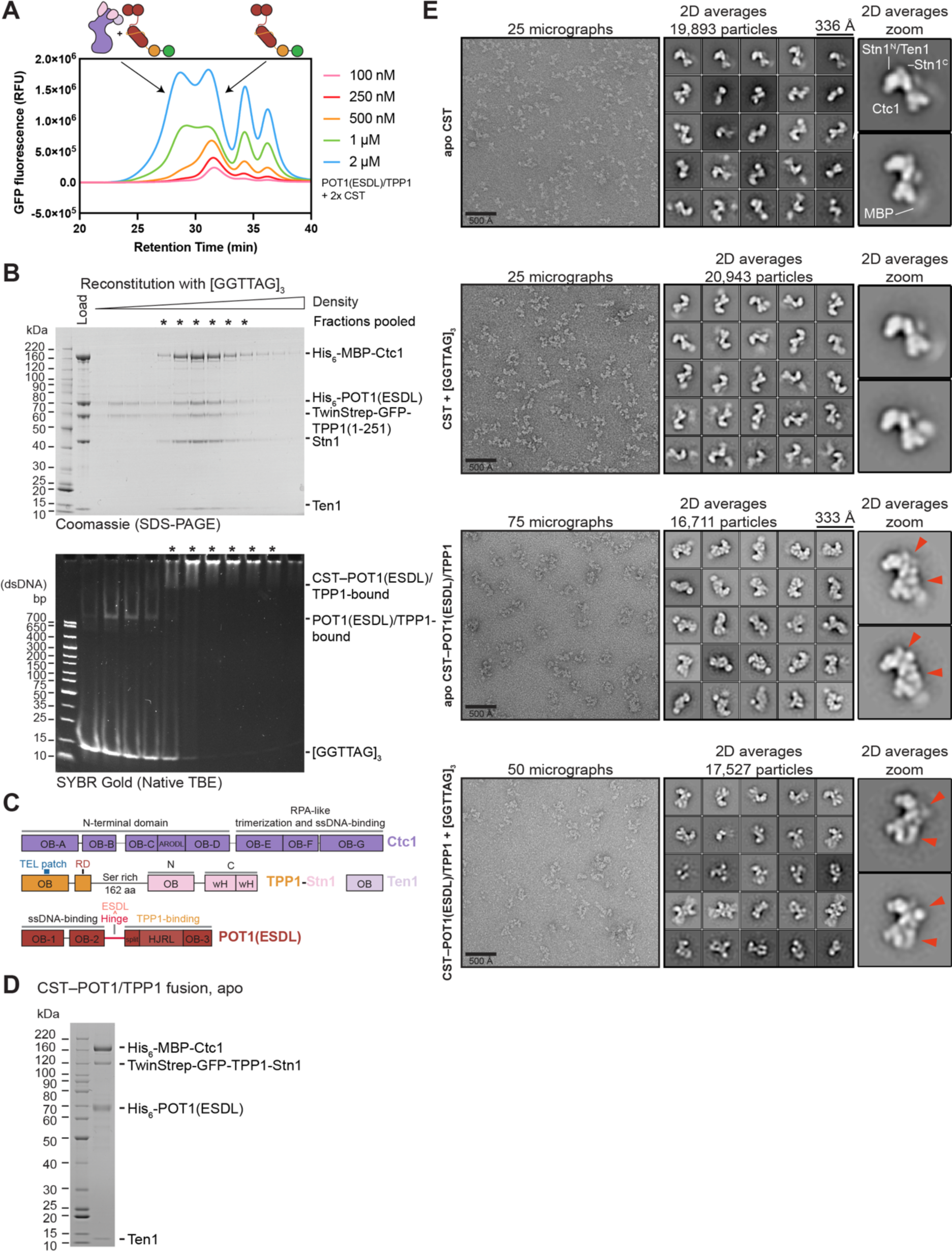
Reconstitution of CST–POT1(ESDL)/TPP1 complexes. (**A**) FSEC analysis of the binding between increasing concentrations of CST and POT1(ESDL)/TPP1 in the absence of telomeric ssDNA showing the concentration dependence of the interaction. RFU: relative fluorescence units. (**B**) Native glycerol gradient analysis of ssDNA-bound CST–POT1(ESDL)/TPP1. Pooled fractions are indicated with an asterisk. (Top) Coomassie-stained SDS-PAGE gel (4-12% Bis-Tris gel run in MOPS-SDS buffer, Invitrogen) of glycerol gradient fractions. (Bottom) SYBR Gold-stained native PAGE (4-20% TBE gel run in 0.5x TB buffer, Invitrogen). (**C**) Domain architecture of components used to reconstitute an apo CST– POT1(ESDL)/TPP1 complex. TPP1 is fused to the N-terminus of Stn1 and retains part of the TPP1 serine-rich linker. (**D**) Coomassie-stained SDS-PAGE gel (4-12% Bis-Tris gel run in MOPS-SDS buffer, Invitrogen) showing the purified CST–POT1(ESDL)/TPP1 fusion complex used for structural analysis. (**E**) Negative-stain EM analysis of CST and CST–POT1(ESDL)/TPP1 complexes. (Left) Representative negative-stain EM micrographs. (Middle) Top 25 reference-free 2D-class averages (sorted by number of particles per class from most populous class at top left to least populous class at bottom right) of each complex. (Right) Enlarged views of selected 2D averages to show CST features. Additional density attributable to the addition of POT1(ESDL)/TPP1 is indicated with red arrowheads.

**Supplemental Fig. S3.**
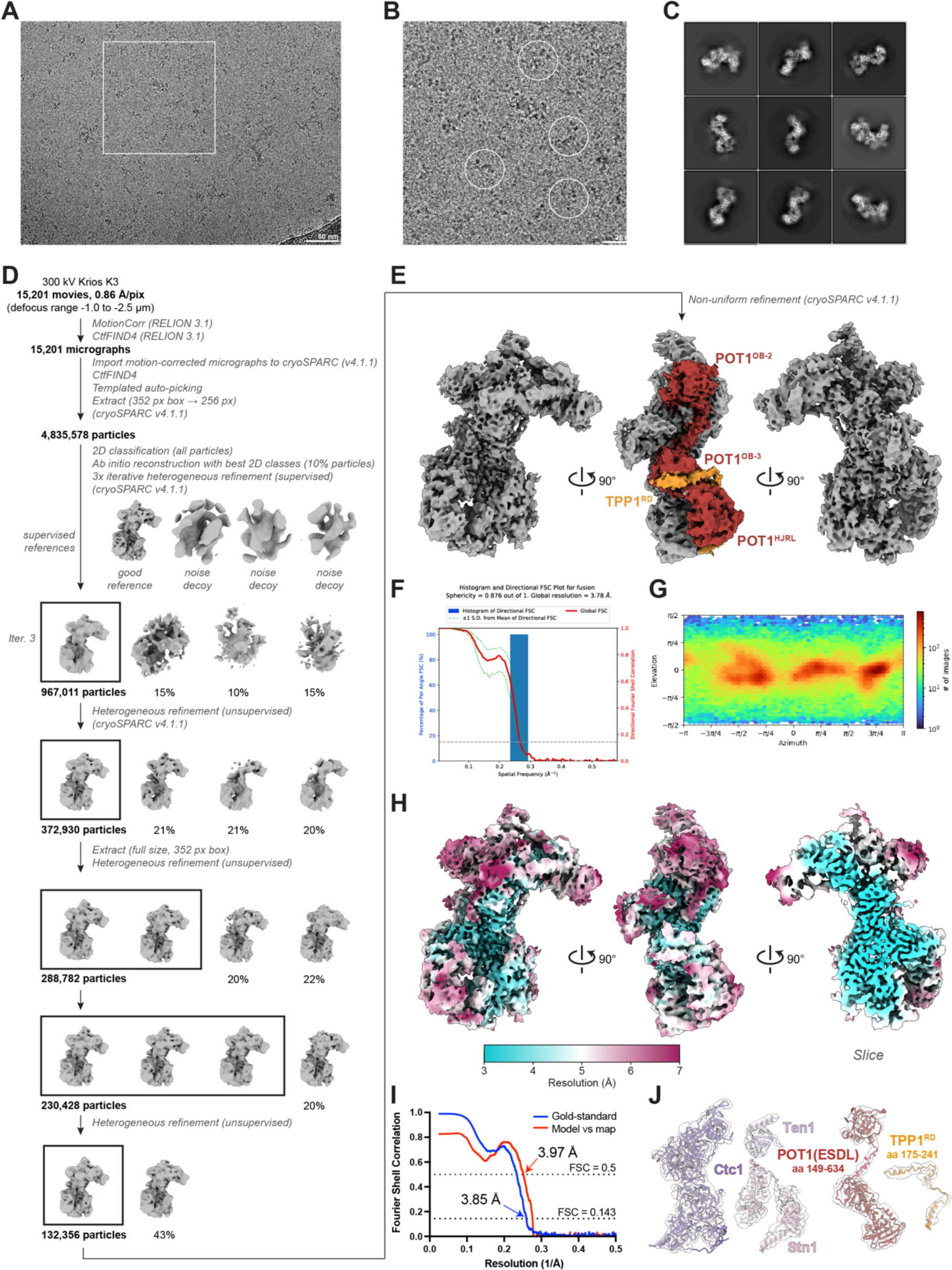
Cryo-EM image processing pipeline for apo CST– POT1(ESDL)/TPP1 complex. (**A**) Representative motion-corrected micrograph. (**B**) Enlarged view of the area marked in (**A**) with selected particles circled. (**C**) Representative 2D-class averages show high-resolution features and different orientations. (**D**) Cryo-EM image-processing pipeline used for the apo CST–POT1(ESDL)/TPP1 complex, including supervised 3D classification with noise decoy classes. Several rounds of heterogeneous refinement were used to select for particles with well-resolved Stn1^C^ and POT1^OB1/2^ (Table 1). (**E**) Final map of the apo CST–POT1(ESDL)/TPP1 complex with POT1(ESDL)/TPP1 colored as in Fig. 2A for reference. (**F**) Directional FSC plots and sphericity values ^65^ of the reconstruction calculated using the 3D-FSC server (https://3dfsc.salk.edu/). (**G**) Plot of the angular distribution of particles in the final reconstruction. (**H**) Local resolution estimates of the apo CST–POT1(ESDL)/TPP1 map. (**I**) Gold-standard (blue) and model-vs-map (red) FSC curves for the apo CST– POT1(ESDL)/TPP1 reconstruction. The map resolution was estimated using the gold-standard FSC and a cut-off criterion of 0.143. The similarity between the resolution estimate of the gold-standard FSC (0.143 cut-off) and resolution estimate of the model-vs-map FSC (0.5 cut-off) suggests no substantial over-fitting. (**J**) Cryo-EM map densities for each subunit indicating quality of fit for the apo CST– POT1(ESDL)/TPP1 model.

**Supplemental Fig. S4.**
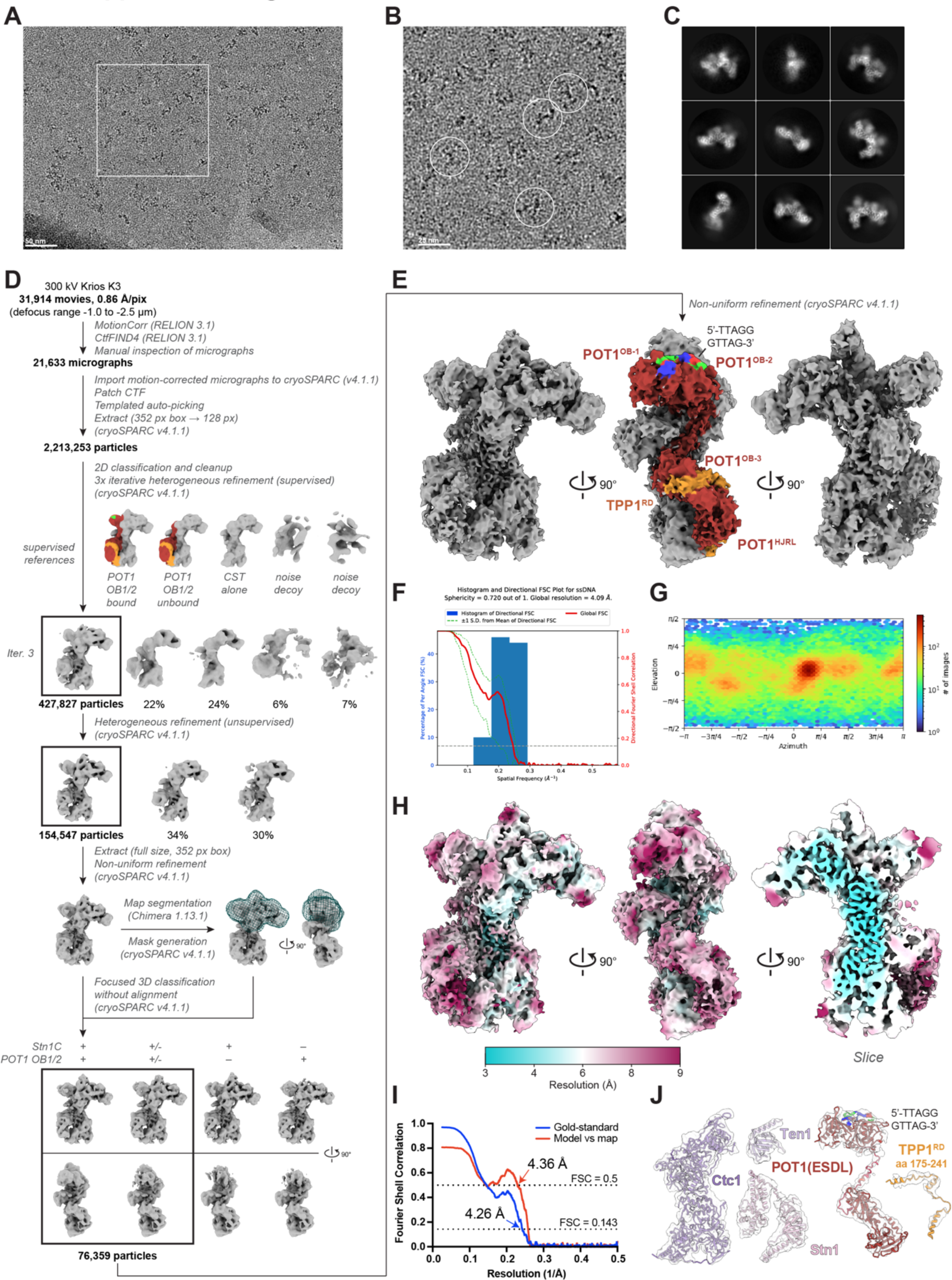
Cryo-EM image processing pipeline for ssDNA-bound CST– POT1(ESDL)/TPP1 complex. (**A**) Representative motion-corrected micrograph. (**B**) Enlarged view of the area marked in (**A**) with selected particles circled. (**C**) Representative 2D-class averages show high-resolution features and different orientations. (**D**) Cryo-EM image-processing pipeline used for the CST–POT1(ESDL)/TPP1–ssDNA complex, including supervised 3D classification with noise decoy classes. Focused 3D classification with a mask was used to select for particles with well-resolved Stn1^C^ and POT1^OB1/2^ (Table 1). (**E**) Final map of the CST–POT1(ESDL)/TPP1–ssDNA complex with ssDNA– POT1(ESDL)/TPP1 colored as in Fig. 2A for reference. (**F**) Directional FSC plots and sphericity values ^65^ of the reconstruction calculated using the 3D-FSC server (https://3dfsc.salk.edu/). (**G**) Plot of the angular distribution of particles in the final reconstruction. (**H**) Local resolution estimates of the CST–POT1(ESDL)/TPP1–ssDNA map. (**I**) Gold-standard (blue), model-vs-map (red) FSC curves for CST–POT1(ESDL)/TPP1– ssDNA reconstruction. The map resolution was estimated using the gold-standard FSC and a cut-off criterion of 0.143. The similarity between the resolution estimate of the gold-standard FSC (0.143 cut-off) and resolution estimate of the model-vs-map FSC (0.5 cut-off) suggests no substantial over-fitting. (**J**) Cryo-EM map densities for each subunit indicating quality of fit for the CST– POT1(ESDL)/TPP1–ssDNA model.

**Supplemental Fig. S5.**
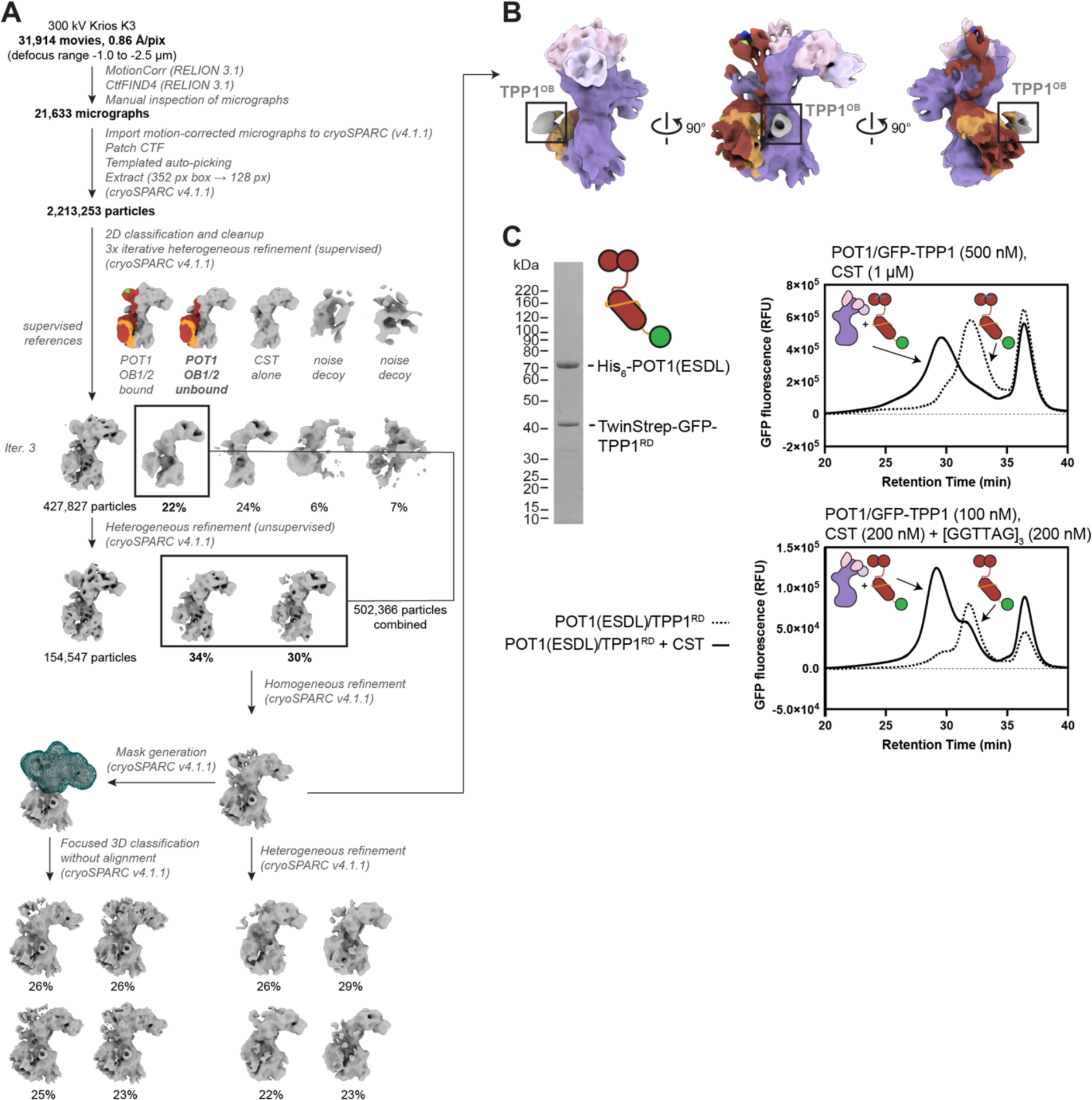
Further cryo-EM image processing of the ssDNA-bound CST–POT1(ESDL)/TPP1 complex and TPP1 OB-fold analysis. (**A**) Alternative cryo-EM image-processing pipeline used for the ssDNA-bound CST– POT1(ESDL)/TPP1 complex. This pipeline was used to select for classes with CST bound to ssDNA that also contained POT1^OB-C^. Briefly, classes matching the supervised reference of this conformation were pooled, but further 3D classification or heterogenous refinement did not yield a high-resolution map, presumably due to remaining conformational heterogeneity. (**B**) When the particles without clear density for POT1^OB-1/2^ and Stn1^C^ were processed, a density representing the TPP1 OB-fold could be seen at low contouring thresholds. Coloring the map of an intermediate step shows an unaccounted-for cylindrical density reminiscent of an OB fold. (**C**) Deletion of the TPP1 OB-fold does not affect the CST–POT1(ESDL)/TPP1 interaction. Protein used and FSEC analysis of CST–POT1(ESDL)/TPP1(ΔOB) interaction in the absence (top) and presence (bottom) of telomeric ssDNA. Traces without CST are shown as dashed lines. RFU: relative fluorescence units.

**Supplemental Fig. S6.**
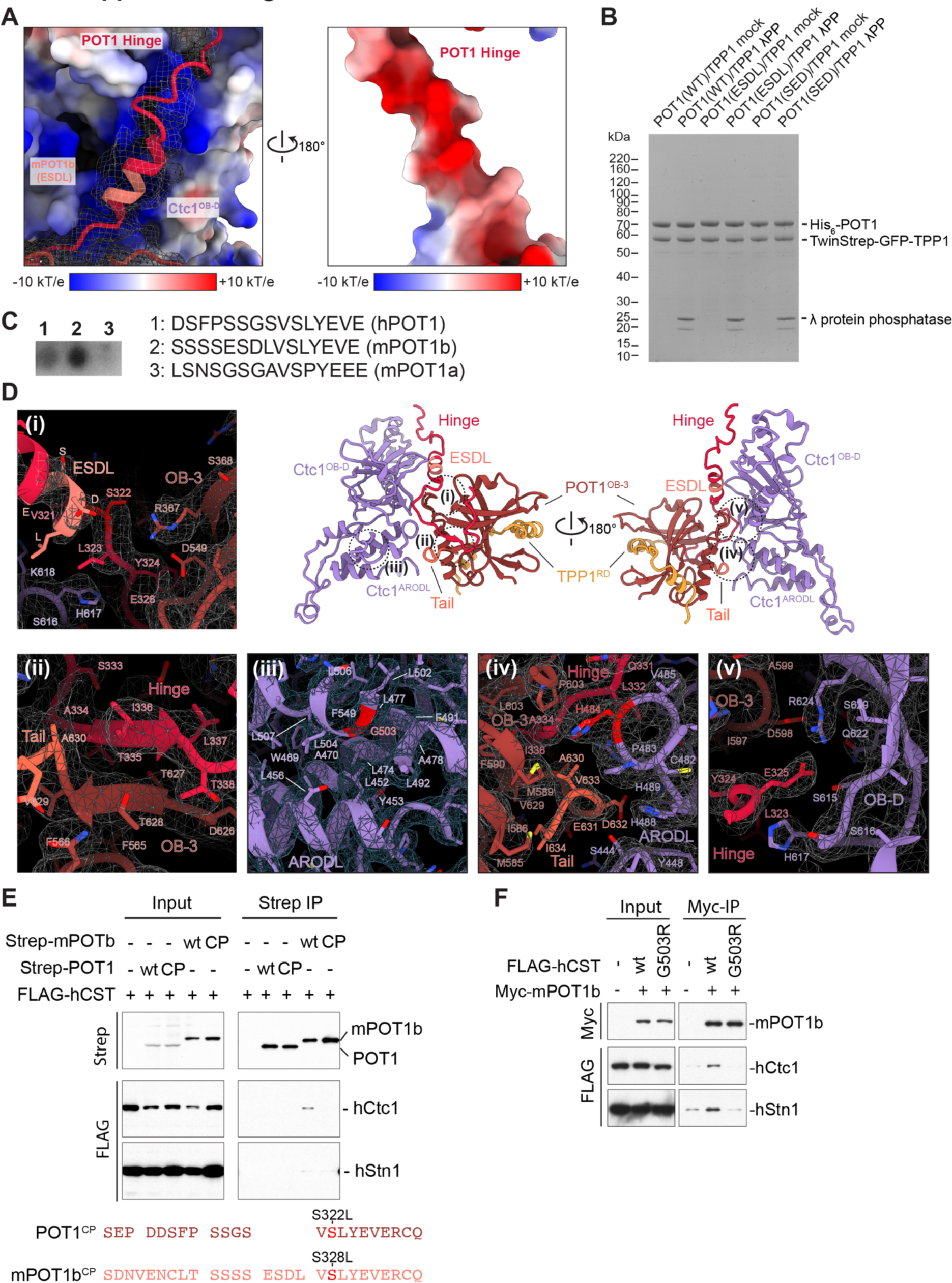
Analysis of the POT1– Ctc1 interaction. (**A**) Electrostatic surface from Fig. 3A with map density of hinge shown as mesh. (Right) 180° rotation view of POT1(ESDL) hinge showing negative charge on the Ctc1-facing side and map density. (**B**) Coomassie blue-stained SDS-PAGE gel (4-12% Bis-Tris gel run in MOPS-SDS buffer, Invitrogen) showing proteins used in the FSEC analysis in Fig. 3C. (**C**) Kinase assay with HeLa nuclear extract on a peptide-scanning array containing peptides corresponding to POT1, mPOT1b, and mPOT1a hinge regions. (**D**) As in Fig. 4B but individual panels include the cryo-EM map density for the apo CST– POT1(ESDL)/TPP1 complex shown as mesh. (**E**) Immunoblots of anti-Strep co-IPs of Strep-tagged POT1 constructs and FLAG-tagged CST from co-transfected 293T cells showing that the corresponding S328L CP mutation in mPOT1b disrupts the CST interaction. Immunoblots were probed with anti-Strep and anti-FLAG antibodies. (**F**) Immunoblots of anti-Myc co-IPs of Myc-tagged mPOT1b and FLAG-tagged CST (wild type or bearing the G503R mutation) from co-transfected 293T cells showing that the CST G503R CP mutation disrupts the mPOT1b interaction. Immunoblots were probed with anti-Myc and anti-FLAG antibodies.

**Supplemental Fig. S7.**
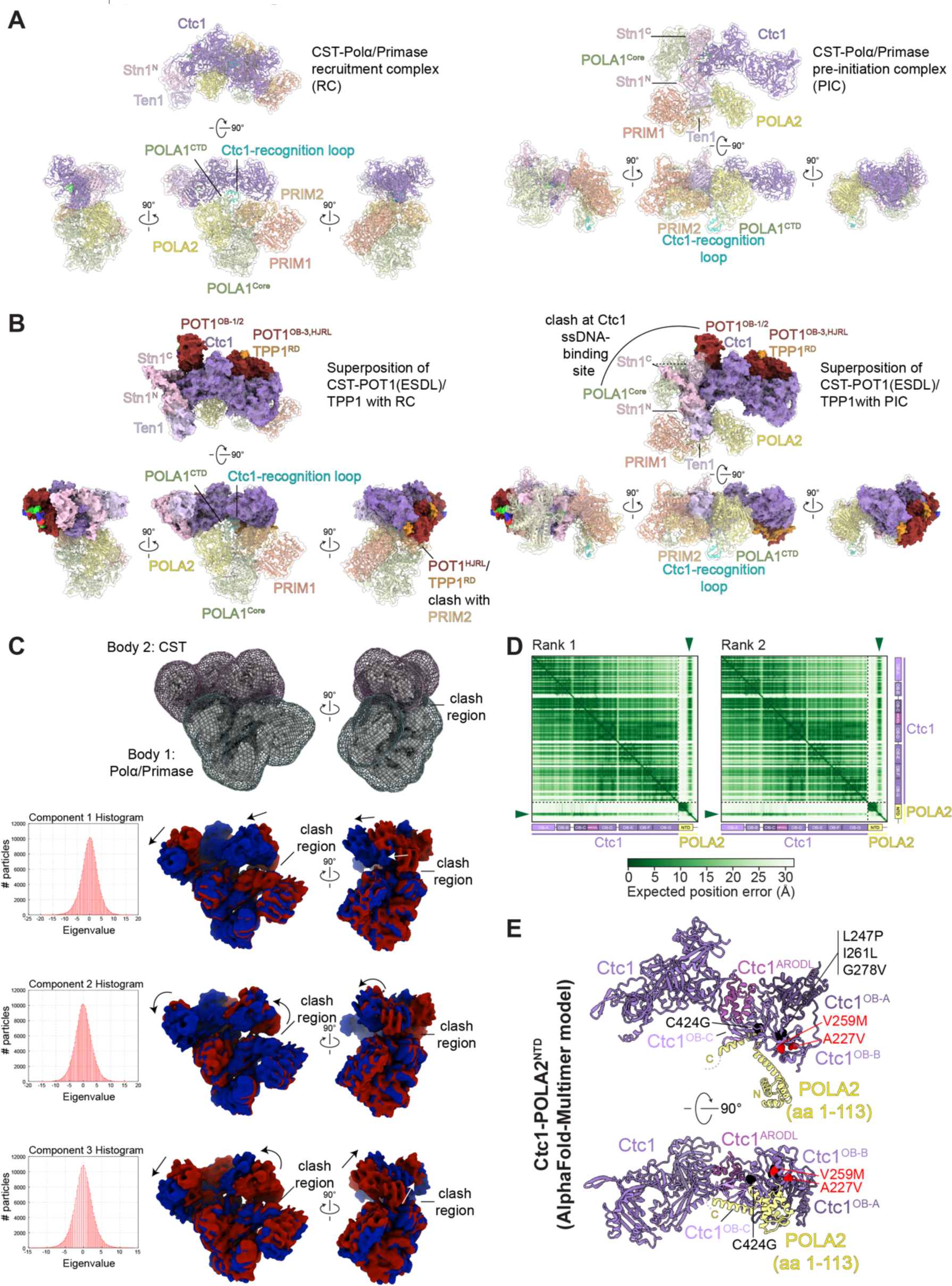
Comparison of the CST–POT1(ESDL)/TPP1 structure with those of CST-Polα/Primase. (**A**) Multiple views of CST–Polα/Primase in RC (left) and PIC (right) conformations. CST– Polα/Primase structures are shown in cartoon representation with a transparent surface. (**B**) Superposition of the CST–POT1(ESDL)/TPP1–ssDNA structure with the structures of the RC (left, see also Video S3) and PIC (right, see also Video S2) complexes showing additional views compared to Fig. 5C. The clash between the POT1^HJRL^ and PRIM2 is indicated. (**C**) Multi-body analysis of CST–Polα/Primase in the RC conformation. Polα/Primase was designated body 1 and CST was designated body 2 with the corresponding masks shown. Histograms of the projections of the relative orientations onto the corresponding components show a unimodal distribution, consistent with continuous flexibility rather than discrete states. The first three principal components account for 61% of the variance in the data. Reconstructed maps from the extreme ends are shown in red and blue for each of the first three principal components with arrows indicating the direction of motion (see also Videos S4-6). (**D**) PAE plots of the top two (of 5) ranked AlphaFold-Multimer ^40^ models for Ctc1–POLA2. Green arrowheads indicate high confidence in the position prediction of POLA2 relative to Ctc1. All 5 top-ranked models predicted the interaction with high confidence. (**E**) AlphaFold-Multimer ^40^ model of Ctc1 bound to POLA2^NTD^. Residues with known CP mutations are shown as spheres. Red spheres indicate mutations previously shown to disrupt Polα/Primase association ^49^; black spheres indicate mutations with unknown mechanism.

## Supplementary Materials

**Supplementary Table S1.** DNA constructs used in this study.

| # | Purpose | Protein | Residues | Vector | Tags |
| --- | --- | --- | --- | --- | --- |
| 1 | Co-IP | hCtc1 | 1-1217 | pLPC | FLAG |
| 2 | Co-IP | hStn1 | 1-368 | pLPC | FLAG |
| 3 | Co-IP | hTen1 | 1-123 | pLPC | FLAG |
| 4 | Co-IP | hTPP1 | 1-458 | pLPC | Myc |
| 5 | Co-IP | hTPP1 Δ10 | 1-448 | pLPC | Myc |
| 6 | Co-IP | hTPP1 Δ20 | 1-438 | pLPC | Myc |
| 7 | Co-IP | hTPP1 Δ30 | 1-428 | pLPC | Myc |
| 8 | Co-IP | hTPP1 Δ40 | 1-418 | pLPC | Myc |
| 9 | Co-IP | hTIN2 | 1-354 (isoform 2) | pcDNA3 | HA |
| 10 | Co-IP | mPOT1b | 1-638 | pLPC | Myc |
| 11 | Co-IP | hPOT1 | 1-634 | pLPC | Myc |
| 12 | Co-IP | HMb1 | hPOT1 1-298 + mPOT1b 300-638 | pLPC | Myc |
| 13 | Co-IP | HMb2 | hPOT1 1-316 + mPOT1b 319-638 | pLPC | Myc |
| 14 | Co-IP | HMb3 | hPOT1 1-320 + mPOT1b 323-638 | pLPC | Myc |
| 15 | Co-IP | HMb4 | hPOT1 1-344 + mPOT1b 350-638 | pLPC | Myc |
| 16 | Co-IP | HMb5 | hPOT1 1-320 + mPOT1b 323-326 +<br>hPOT1 321-344 + mPOT1b 350-638 | pLPC | Myc |
| 17 | Co-IP | EE1 | hPOT1 1-634 (S317E/S322E) | pLPC | Myc |
| 18 | Co-IP | EE2 | hPOT1 1-634 (S317E/S318E) | pLPC | Myc |
| 19 | Co-IP | ED | hPOT1 1-634 (S317E/G319D) | pLPC | Myc |
| 20 | Co-IP | EED | hPOT1 1-634<br>(S317E/S318E/G319D) | pLPC | Myc |
| 21 | Co-IP | SED | hPOT1 1-634<br>(S315E/S317E/G319D) | pLPC | Myc |
| 22 | Co-IP | hPOT1 | 1-634 | pQE | Strep |
| 23 | Co-IP | hPOT1<sup>CP</sup> | 1-634 (S322L) | pQE | Strep |
| 24 | Co-IP | mPOT1b | 1-638 | pQE | Strep |
| 25 | Co-IP | mPOT1b<sup>CP</sup> | 1-638 (S328L) | pQE | Strep |
| 26 | Co-IP | hCtc1<sup>G503R</sup> | 1-1217 (G503R) | pQE | FLAG |
| 27 | Protein expression | CST | [Ctc1 1-1217]<br>[Stn1 1-368]<br>[Ten1 1-123] | pBIG1a | His<sub>6</sub>-MBP-3C |
| 28 | Protein expression | POT1(WT)/<br>TPP1 | [POT1 1-634]<br>[TPP1 1-251] | pBIG1b | His<sub>6</sub>-<br>TwinStrep-GFP-TEV |
| 29 | Protein expression | POT1(ESDL)/<br>TPP1 | [POT1(ESDL) 1-638]<br>[TPP1 1-251] | pBIG1b | His<sub>6</sub>-<br>TwinStrep-GFP-TEV |
| 30 | Protein expression | POT1(SED)/<br>TPP1 | [POT1 (SED) 1-634]<br>[TPP1 1-251] | pBIG1b | His<sub>6</sub>-<br>TwinStrep-GFP-TEV |
| 31 | Protein expression | POT1(ESDL)<br>TPP1(ΔOB) | [POT1(ESDL) 1-638]<br>[TPP1 159-251] | pBIG1b | His<sub>6</sub>-<br>TwinStrep-GFP-TEV |
| 32 | Protein expression | Fusion CST–<br>POT1/TPP1<br>complex | [Ctc1 1-1217]<br>[TPP1 (1-397)–GGS–<br>GGS–Stn1 (1-368)]<br>[Ten1 1-123]<br>[POT1(ESDL) 1-638] | pBIG2ab | His<sub>6</sub>-MBP-3C<br>TwinStrep-<br>GFP-TEV<br>His<sub>6</sub>- |

**Supplementary Video S1. Cryo-EM map and model of CST–POT1(ESDL)/TPP1 complexes.** 360° rotation of the cryo-EM map and model from apo and ssDNA-bound CST–POT1(ESDL)/TPP1. Colors are the same as in Fig. 2.

**Supplementary Video S2. Superposition of ssDNA-bound CST–POT1(ESDL)/TPP1 with CST–Polα/Primase pre-initiation complex.** Rotation movie showing superposition of CST–POT1(ESDL)/TPP1–ssDNA complex (surface representation; solid colors as in Fig. 2) with CST–Polα/Primase PIC (surface representation at 50% opacity; cartoon model shown underneath). Colors of Polα/Primase are the same as in ^18^. PRIM1-salmon; PRIM2-light orange; POLΑ1-light green (Ctc1-recognition loop-cyan); POLA2-yellow.

**Supplementary Video S3. Superposition of ssDNA-bound CST–POT1(ESDL)/TPP1 with CST–Polα/Primase recruitment complex.** Rotation movie showing superposition of CST–POT1(ESDL)/TPP1–ssDNA complex (surface representation; solid colors as in Fig. 2) with CST–Polα/Primase RC (surface representation at 50% opacity; cartoon model shown underneath). Colors of Polα/Primase are the same as in ^18^. PRIM1-salmon; PRIM2-light orange; POLΑ1-light green (Ctc1-recognition loop-cyan); POLA2-yellow.

**Supplementary Videos S4-6. Multi-body refinement movies of the first three principal motions contributing to CST-Polα/Primase RC flexibility.** Components 1, 2, and 3, respectively.

## Notes

### Summary of Updates

This version of the manuscript has been revised with an expanded introduction and discussion, and the figures have been slightly reformatted.

